# Sex and Gender Considerations in Reporting Guidelines of Health Research: A Systematic Review

**DOI:** 10.1101/2021.04.06.438690

**Authors:** Amédé Gogovor, Hervé Tchala Vignon Zomahoun, Giraud Ekanmian, Évèhouénou Lionel Adisso, Alèxe Deom Tardif, Lobna Khadhraoui, Nathalie Rheault, David Moher, France Légaré

## Abstract

**Background:** Despite growing recognition of the importance of sex and gender considerations in health research, they are rarely integrated into research design and reporting. We sought to assess the integration of sex, as a biological attribute and gender as a socially constructed identity in published reporting guidelines.

**Methods and Findings:** We conducted a systematic review of published reporting guidelines listed on the EQUATOR website (www.equator-nework.org) from inception until December 2018. We selected all reporting guidelines (original and extensions) listed on the EQUATOR library. We used EndNote Citation Software to build a database of the statement of each guideline identified as ‘full bibliographic reference’ and retrieved the full texts. Reviewers independently extracted the data from the checklist/abstract/main text of guidelines. Data were analyzed using descriptive statistics and narrative synthesis. A total of 407 reporting guidelines were included; they were published between 1995 and 2018. Of the 407 guidelines, 159 (39%) mentioned “sex” and/or “gender” in the checklist/abstract/main text. Of these, 90 (22.1%) mentioned only “sex”, and 91 (22.4%) mentioned only “gender”. In the checklist of the reporting guidelines (n = 363), “sex” and “gender” were mentioned in 50 (13.8%) and 39 (10.7%), respectively. Only one reporting guideline met the three criteria of correct use of sex and gender concepts. Trends in the use of sex and gender in the checklists showed that the use of “sex” only started in 2003, while “gender” has been used since 1996.

**Conclusions:** We assessed the integration of sex and gender considerations in reporting guidelines based on the use of sex- and gender-related words. Our findings showed a low use and integration of sex and gender concepts in reporting guidelines. Authors of reporting guidelines should reduce this gap for a better use of research knowledge.

**Registration:** PROSPERO no. CRD42019136491.

## INTRODUCTION

Deficiencies in the quality of reporting of health research are well documented in the literature [1, 2]. Consequences of inadequate reporting include lapses of scientific integrity and difficulty in judging the reliability of the results and the relevance of the evidence [2].

One of the recurrent deficiencies in research design and reporting is the lack of the integration of sex and gender considerations. Despite growing recognition of the importance of sex and gender in the manifestation and management of health conditions, their considerations are rarely integrated in research design and reporting [3–5]. This limitation may further explain why there is waste in research, as research being performed right now is not aligned with or does not reflect the sex and gender profiles of the population.

Based on the knowledge-to-action process [6], synthesizing gaps in the integration of sex and gender and developing appropriate reporting guidelines will contribute to effective knowledge translation. Sex refers to “a set of biological attributes in humans and animals and is primarily associated with physical and physiological features, including chromosomes, gene expression, hormone levels and function, and reproductive/sexual anatomy”[7]. The traditional categorization of sex is dichotomous as male or female; sometimes, other is offered as a response option. Gender refers to “the socially constructed roles, behaviours, expressions and identities of girls, women, boys, men, and gender diverse people and it influences how people perceive themselves and each other, how they act and interact, and the distribution of power and resources in society”[7]. Categories of gender include men, women, and gender-diverse people. Thus, “sex” and “gender” hold different meanings and should not be used interchangeably [8–10].

Reporting guidelines are developed to improve the transparency, accuracy and completeness of reporting for different types of research [11, 12]; thus, these guidelines need to incorporate items related to sex and gender considerations to provide comprehensive guidance. A reporting guideline is defined as “a checklist, flow diagram, or explicit text to guide authors in reporting a specific type of research, developed using explicit methodology”[2]. A recent study examined the inclusion of sex and gender considerations in the publishing guidelines of several top-ranking health journals, and the study made recommendations to strengthen these considerations in policies and practices of health journals [13]. However, no systematic investigation of sex and gender considerations in reporting guidelines has been made. The aim of this study was to assess the integration of sex and gender concepts in published reporting guidelines based on the use of sex- and gender-related words. We examined the correct use of sex and gender concepts, the publication trends in the use of sex and gender terms, and the nature of sex and gender information in the checklist.

## METHODS

The protocol of the systematic review is registered with the International Prospective Register of Systematic Reviews (PROSPERO CRD42019136491) [14]. We followed the Preferred Reporting Items for Systematic Reviews and Meta-Analyses (PRISMA) statement [15](S1 Checklist) to guide the report.

### Eligibility criteria, information sources and study selection

The EQUATOR Network Team, which maintains a collection of reporting guidelines for health research, has developed search strategies for PubMed, Embase, Cinahl, and Web of Science to identify reporting guidelines published since 1996 in English. Free-text and controlled vocabulary terms used in the search strategies include reporting guideline(s); reporting standard(s); reporting guidance; reporting requirement(s); reporting criteria; reporting recommendation(s); reporting checklist(s); reporting statement and reporting instruction(s). The search strategies are run regularly [16], and the results are systematically reviewed by the EQUATOR Network Team for inclusion in the database of reporting guidelines.

We systematically included all published and listed reporting guidelines (original and extensions) in the Equator Network registry (www.equator-nework.org), as of 31 December 2018; thus, the selection flowchart was not applicable. We used EndNote (EndNote Citation Software, Version 9.3, Clarivate Analytics, New York, NY, USA) to build a database of the statement of each reporting guideline identified as a “full bibliographic reference” on EQUATOR and retrieved the full texts. This allowed us to be consistent by having one main document (statement) per reporting guideline and was justified by the fact that authors rarely consult explanation and elaboration documents [17]. For reported guidelines that included sex and gender terms in their checklist, we consulted complementary documents listed on their webpage in the EQUATOR Network Registry to complete the assessment of correct use of these terms.

### Data collection process

Pairs of reviewers (AG, GE, ÉLA, ADT) independently extracted study data using a pretested extraction form. The information extracted included the following: 1) ***Characteristics of the reporting guidelines*** (e.g., author, year, title, acronym, type of study as documented on EQUATOR (randomized trial, observational, systematic review, protocol, diagnostic/prognostic, case report, clinical practice guideline, qualitative research, animal preclinical, quality improvement, economic evaluation, experimental, other, and ‘nonspecific’ for reporting guidelines that do not apply to any specific type of study)); 2) ***Description of the integration of sex and gender*** (e.g., presence of any of sex- and gender-related words (sex, gender, female(s), male(s), man, woman, men, women, boy(s), girl(s), gender-diverse) in the checklist, flowchart and main text and the number of occurrences of these words in each using electronic text search tool. For scanned documents and images, we used a free online Optical Character Recognition (OCR) tool to convert them into editable text. The main text (statement) includes text from introduction to conclusion. We also extracted sex- and gender-related words from the abstract and the reference list of reporting guidelines included. Sex and gender related words in the following sections were excluded: affiliation, acknowledgment, tables, and figures. 3) ***Correct use of sex or gender*** based on established definitions of sex and gender [7, 18]. The correct use of sex and gender was assessed by three criteria: nonbinary use, the use of appropriate categories, and noninterchangeable use of sex and gender. The three criteria are described elsewhere [19] and summarized in table 1. The use of sex and gender was correct if all three criteria were met, incorrect if at least one of three criteria was not met, and unclear if at least one of the criteria was reported as unclear and the others were met. We consulted explanation and elaboration documents of reported guidelines for the assessment of correct use of sex and gender concepts in their checklist. 4) Type of sex and gender information in the checklist. Additionally, a sample of 10% of the reporting guidelines was searched manually and reported (S1 Image) for comparison with the electronic search in the following sections: checklist, abstract, statement, and references. We contacted the author of one reporting guideline and obtained a copy of the checklist because a supplemental document was not available. Discrepancies were resolved by consensus between two team members, and with a third member when necessary. We performed double counting to resolve discrepancies between electronic versus manual searches.

**Table 1.** Criteria for assessing correct use of sex and gender terms in reporting guidelines

| Criterion | Definition |
| --- | --- |
| **Nonbinary** | 1. Nonbinary use: male, female or intersex for sex; man (men), woman (women) or gender-diverse for gender; description of sex or gender implies more than two categories<br>2. Binary-use: male or female for sex; boy/man/men or girl/woman/women for gender; description of sex or gender |
|  | implies two categories<br>3. Unclear: terms of sex or gender used without specification of categories |
| **Appropriate categories** | 1. Appropriate: consistent use of male/female/intersex for sex; boy/man/men, girl/woman/women or gender-diverse for gender<br>2. Inappropriate: inconsistent use of male/female/intersex for sex; boy/man/men, girl/woman/women or gender-diverse for gender; e.g., male/female/intersex for gender<br>3. Unclear: terms of sex or gender used without specification of categories |
| **Noninterchangeability** | 1. Noninterchangeable: consistent use of sex to describe biological attributes and gender for sociocultural attributes<br>2. Interchangeable: inconsistent use of sex to describe biological attributes and gender for sociocultural attributes; e.g., indiscriminate use of sex and gender, male and man for the same concept<br>3. Unclear: terms of sex or gender used without specification of categories, any other situation where assessment is unrealizable |

Adapted from Adisso et al[19]

### Internal validity assessment

The assessment of internal validity ensures the minimization of potential bias in the development of reporting guideline recommendations. To assess the quality of the included reporting guidelines, we adapted the risk of bias checklist developed by Cukier et al. [20] since there is no validated tool for internal validity assessment for methods systematic reviews. One author (AG) first drafted a list of the three criteria relevant to reporting guideline development [2]. The list of items was reviewed by an author (DM) with extensive experience in the development of reporting and pilot tested (two rounds) by members of the author group for consistency and feasibility. The final 3-item internal validity assessment checklist is shown in box 1 (detailed coding manual in supplementary file S1 manual). Each item was coded “yes”, “no”, or “unclear”. The judgment rule to determine evidence-based development of the included reporting guidelines was high internal validity (i.e., low risk of bias) if ≥ 2 “yes”. The assessment was based on the main text (statement) of each reporting guideline. For criterion 1, when a specific group was named the developer of the guideline, we consulted the internet to determine whether it represented more than one stakeholder group. Pairs of three reviewers (AG, GE, HTVZ) independently conducted the assessment and any disagreements were resolved by consensus or third-party adjudication.

##### Box 1. List of quality assessment criterion

1. Did the developers of the guideline represent more than one stakeholder group (e.g., researchers, funders, publishers)?
2. Did the developers report gathering any data for the creation of the guideline (e.g., carry out a literature review, collect anecdotal data)?
3. Did the developers report the use of a consensus process (e.g., Delphi, RAND/UCLA Appropriateness Method, nominal group technique, consensus meeting, development conference)?

### Data Synthesis and Analysis

We analyzed extracted data using descriptive statistics and reported the numbers and percentages of reporting guidelines that integrate sex- and/or gender-related items. We also calculated the mean number of occurrences of sex- or gender-related words in the different sections of reported guidelines, and we calculated the mean number of sex and gender terms in the checklists for each year of publication. We reported publication trends over time in the use of sex and gender in the checklists. For the reporting guidelines that integrated sex and/or gender terms in their checklist, we calculated the percentage of reporting guidelines that met each of the three criteria; and the percentage of correct use of sex and gender, i.e., all three criteria met.

We qualitatively synthesized the nature of sex and gender information in the checklist into three groups: i) mention of sex and/or gender terms with no description of the categories; ii) use of sex or gender terms with description of the categories; and iii) detailed definition or description on how sex and/or gender should be integrated. Finally, we compared the frequencies of the use of sex- and gender-related words for electronic and manual modes of assessment. We identified the number of times the terms of interest were used throughout the different sections of reporting guidelines. This identification was done both electronically and manually to ensure consistency of our results. There was concordance if the numbers were the same. SAS (SAS, version 9.4, Institute, Cary, NC, USA) and R (R software, version 3.6, R Core Team, University of Auckland, New Zealand) were used to perform the analyses.

## RESULTS

### Characteristics of the included reporting guidelines

A total of 407 reporting guidelines (statements) were included in the review, published between 1995 and 2018. Seven related explanation and elaboration documents were consulted during the assessment of correct use of “sex” and “gender”. The most prevalent year of publishing reporting guidelines was 2010 with 51 reported guidelines (Figure 1). Of 407 included reporting guidelines, 66 (16.2%) were extensions of existing guidelines, 363 (89.2%) included a checklist and 26 (6.4%) a flowchart. While 349 (85.8%) included an abstract, only 134 (33%) out of 407 reporting guideline statements adopted the traditional structure of a scientific article with the following sections: introduction, methods, results, and conclusion. Most reporting guidelines 243 (59.7%) were developed for specific methodological approaches (for part of/whole report) while 122 (29.9%) were developed for a specific type of study. Based on the classification on the EQUATOR website, the most common study types targeted by the included reported guidelines were ‘experiment’ (145, 35.6%), ‘randomized trials’ (132, 32.4%), and ‘observational’ (115, 28.3%) (Table 2). A reference list of included reporting guidelines is shown in S1 Table.

**Figure 1.**
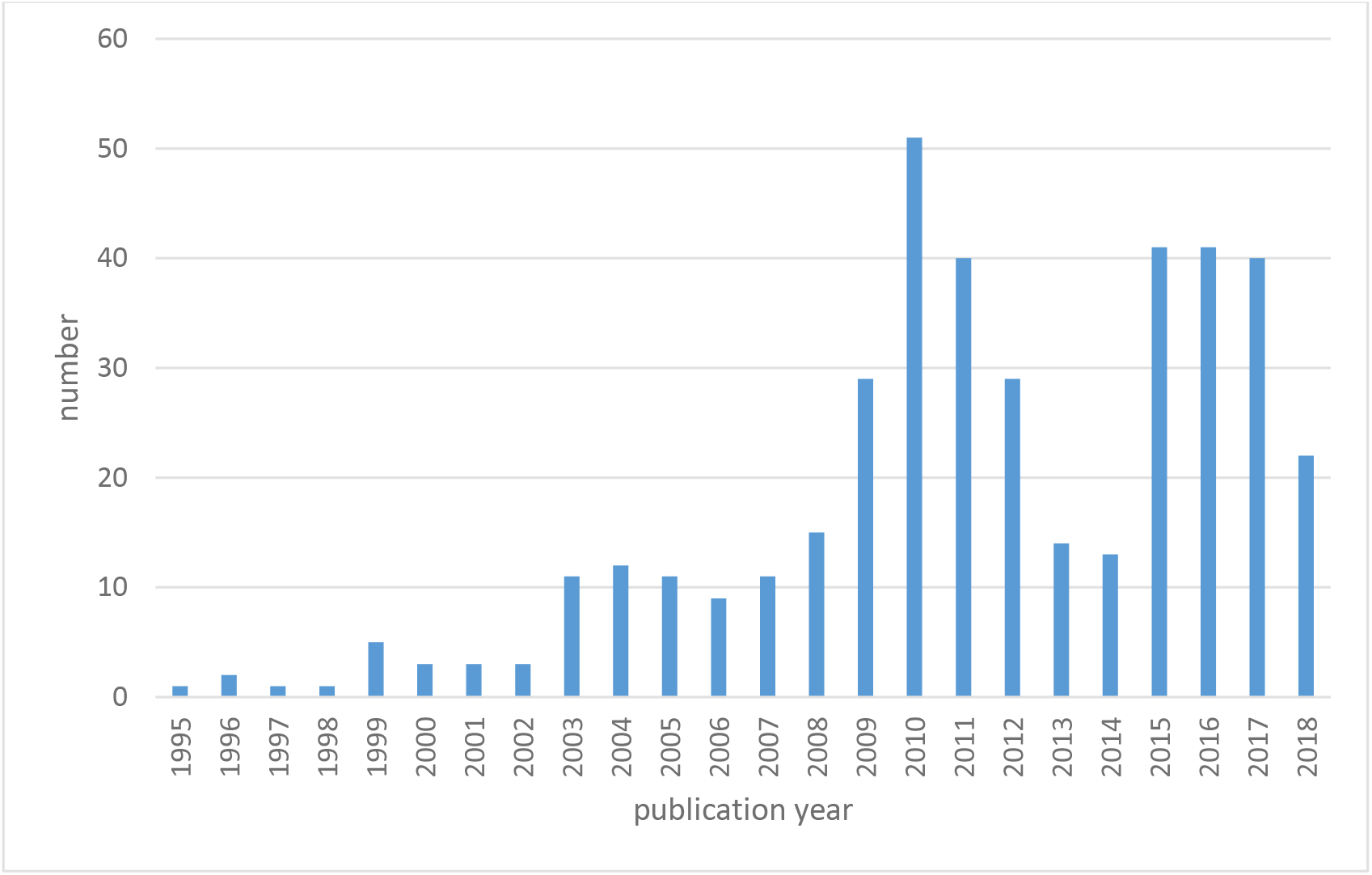
Distribution of reporting guidelines per year

**Table 2.** Distribution of reporting guidelines by study types

| Study type | N (407)<sup>1</sup> | % |
| --- | --- | --- |
| **Case report<sup>2</sup>** | 4 | 1.0 |
| **Clinical practice guideline** | 8 | 2.0 |
| **Diagnostic/prognostic** | 18 | 4.4 |
| **Economic evaluation** | 16 | 3.9 |
| **Experiment** | 145 | 35.6 |
| **Nonspecific<sup>3</sup>** | 103 | 25.3 |
| **Observational** | 115 | 28.3 |
| **Other<sup>4</sup>** | 16 | 3.9 |
| **Preclinical** | 15 | 3.7 |
| **Protocol** | 10 | 2.5 |
| **Quality improvement** | 5 | 1.2 |
| **Qualitative** | 17 | 4.2 |
| **Randomized trial** | 132 | 32.4 |
| **Systematic review** | 34 | 8.4 |

<sup>1</sup>Numbers do not add up to N because reporting guidelines can be classified in more than one category
<sup>2</sup>Based on the category of study types of EQUATOR homepage
<sup>3</sup> Do not apply to any specific type of study
<sup>4</sup>As specified on the individual page of reporting guidelines on EQUATOR

### Electronic versus manual search of words

A total of 41 (10%) reporting guidelines were randomly selected for the comparison of electronic and manual identification of sex and gender-related words. We found 17 discrepancies in 12 reporting guidelines and only one discrepancy was in favor of manual search, after verification (S2 Table). The concordance between the electronic and manual searches of sex, gender and related words was 97.6% for “sex”, “women” and “men” and 100% for “gender”, “female”, “male”, “woman”, “man”, “boy”, and “girl” in the checklists. In the statement, the concordance was 95.1% for “gender”, “women”, and “men”; 97.6% for “male”; and 100% for “sex”, “man”, “female”, “boy” and “girl”. The references section showed a concordance of 95.1% for “sex”; 97.6% for “gender”, “women”, and “men”; and 100% for “woman”, “man”, “female”, “male”, “boy” and “girl”. No discrepancies were found in the abstract section (S3 Table).

### Internal validity assessment

We conducted the assessment on a random number of 100 reported guidelines of 407 included due to time and resource constraints. The summary of the assessment is presented in figure 2 (S1 Database for detailed internal validity assessment). There was evidence-based development of just above the average of the assessed reporting guidelines, i.e., high internal validity (53/100).

**Figure 2.**
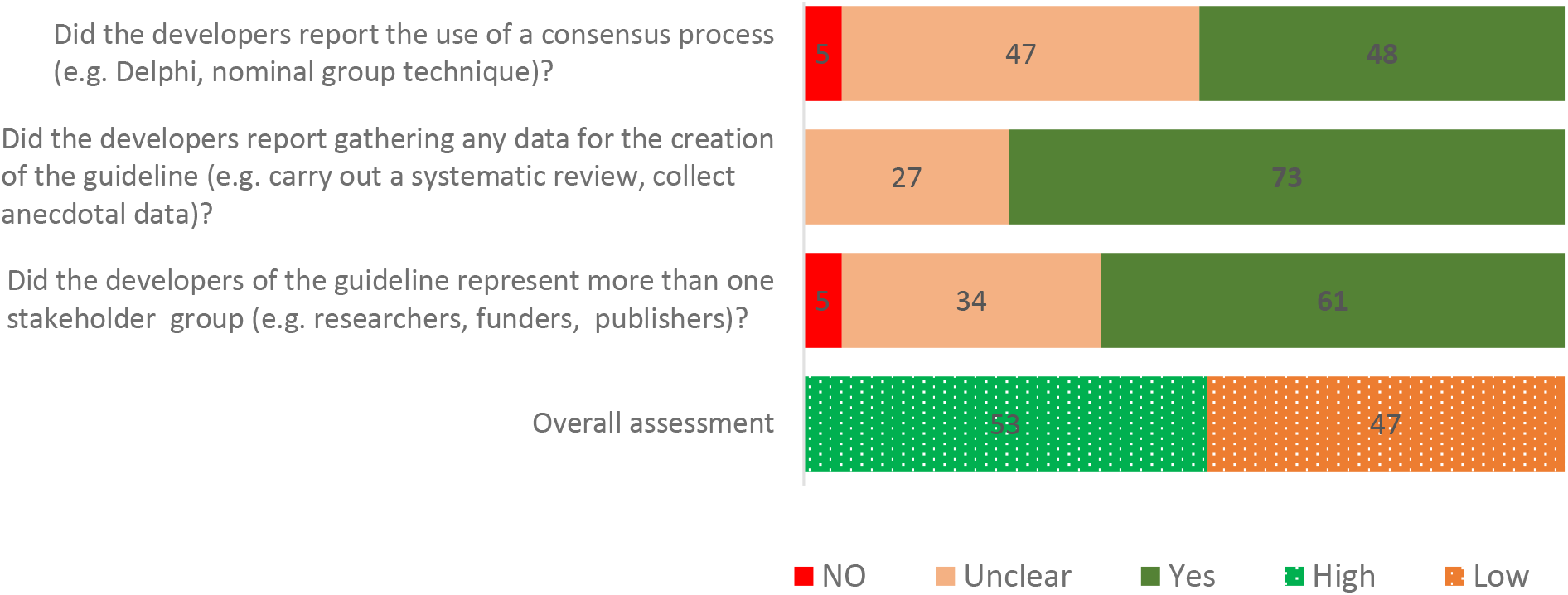
Summary of internal validity assessment

### Description of the integration of sex- and gender-related words

#### Use of sex- and gender-related words

Of 407 reporting guidelines, 233 (57.3%) mentioned at least one of the sex- and gender-related words (sex, gender, female(s), male(s), man, woman, men, women, boy(s), girl(s)). The distribution of the use of sex- and gender-related words in the different sections of reporting guideline statements is shown in table 3. In the checklist of the reporting guidelines (n = 363), only 50 (13.8%) used “sex”, 39 (10.7%) used “gender”, 85 (23.4%) used “sex” and/or gender, and 4 (1%) used both. In the main text (statement), “sex” and “gender” were mentioned in 14.2% and 17.9%, respectively. The most common sex- and gender-related words used in the main text (statement) were “gender” (73/407, 17.9%), “woman/women” (64/407, 15.7%), “sex” (58/407, 14.2%), and “male(s)” (42/407, 10.3%). Distributions of the use of “sex” and “gender” terms according to study types and sections of reported guidelines are presented in supplementary tables S4 and S5.

**Table 3.** Percentage of the presence of sex- and gender-related words in different sections of reporting guidelines

|  | **Checklist**<br>**N = 363** | **Flowchart**<br>**N = 26** | **Abstract**<br>**N = 349** | **Statement**<br>**N = 407** | **References**<br>**N = 407** |
| --- | --- | --- | --- | --- | --- |
| **Sex** | 50 (13.8) | 1 (3.9) | 3 (0.9) | 58 (14.2) | 27 (6.6) |
| **Gender** | 39 (10.7) | 1 (3.9) | 5 (1.4) | 73 (17.9) | 19 (4.7) |
| **Female(s)** | 10 (2.8) | 1 (3.9) | 2 (0.6) | 39 (9.6) | 13 (3.2) |
| **Male(s)** | 9 (2.5) | 1 (3.9) | 2 (0.6) | 42 (10.3) | 15 (3.7) |
| **Man/Men** | 7 (1.9) | 0 (0.0) | 5 (1.4) | 39 (9.6) | 39 (9.4) |
| **Woman/**<br>**Women** | 1 (0.3) | 0 (0.0) | 1 (0.3) | 11 (2.7) | 2 (0.5) |
| **Women** | 6 (1.7) | 1 (3.9) | 4 (1.1) | 64 (15.7) | 65 (15.9) |
| **Boy(s)** | 0 (0.0) | 0 (0.0) | 0 (0.0) | 1 (0.1) | 5 (1.0) |
| **Girl(s)** | 0 (0.0) | 0 (0.0) | 0 (0.0) | 6 (1.5) | 5 (1.2) |
| **No Sex- or**<br>**Gender-**<br>**related words** | 271 (74.7) | 23 (88.5) | 337 (96.6) | 241 (59.2) | 294 (72.2) |

Distributions are reported as n (%). Statement refers to the main text from introduction to discussion/conclusion, excluding tables, figures, acknowledgement, and affiliations. Percentages do not equal 100% because sections can include more than one word.

#### Correct use of sex and gender terms

Table 4 reports the correct use of “sex” and “gender” terms in the reporting guidelines that used sex and gender terms in their checklist. The correct use of “sex” and “gender” terms was assessed in the reporting guidelines that mentioned these terms in their checklist (n=85). The use was correct in only one reporting guideline published in 2016 (all three criteria met) (S1 Table, RG102); incorrect in 22 (25.9%) and unclear in 62 (72.9%). Few reporting guidelines met the individual criteria: 4 (4.7%) for nonbinary use (use of “gender-diverse” in one guideline (S1 Table, RG102), “transgender” in two (S1 Table, RG102, RG188), and other categories in two (S1 Table, RG128, RG263); 5 (5.9%) for noninterchangeable; and 9 (10.6%) for the use of appropriate categories.

**Table 4.**
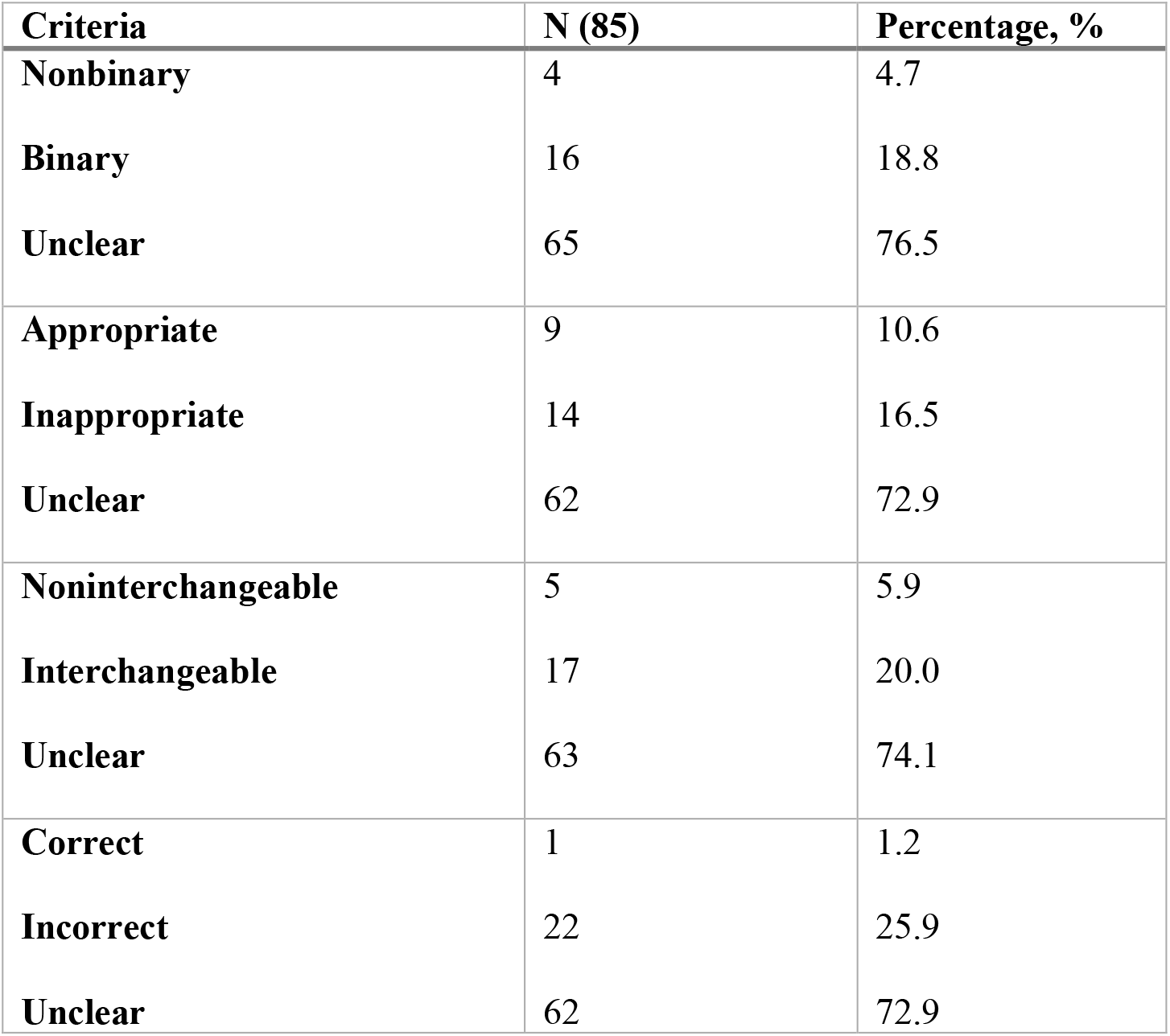
Correct use of sex or gender terms in reporting guidelines that included these terms in their checklist

| **Criteria** | **N (85)** | **Percentage, %** |
| --- | --- | --- |
| **Nonbinary** | 4 | 4.7 |
| **Binary** | 16 | 18.8 |
| **Unclear** | 65 | 76.5 |
| **Appropriate** | 9 | 10.6 |
| **Inappropriate** | 14 | 16.5 |
| **Unclear** | 62 | 72.9 |
| **Noninterchangeable** | 5 | 5.9 |
| **Interchangeable** | 17 | 20.0 |
| **Unclear** | 63 | 74.1 |
| **Correct** | 1 | 1.2 |
| **Incorrect** | 22 | 25.9 |
| **Unclear** | 62 | 72.9 |

#### Trends in the use of sex and gender terms

Publication trends (from 1995 to 2018) in the use of sex and gender terms in checklists showed that “sex” and “gender” were increasingly used over time (figure 3) based on the proportion of reporting guidelines. A similar trend was observed when considering the mean number of “sex” and “gender” occurrences in the checklists (S1 figure). While “gender” has been used since 1996, the use of “sex” started only in 2003.

**Figure 3.**
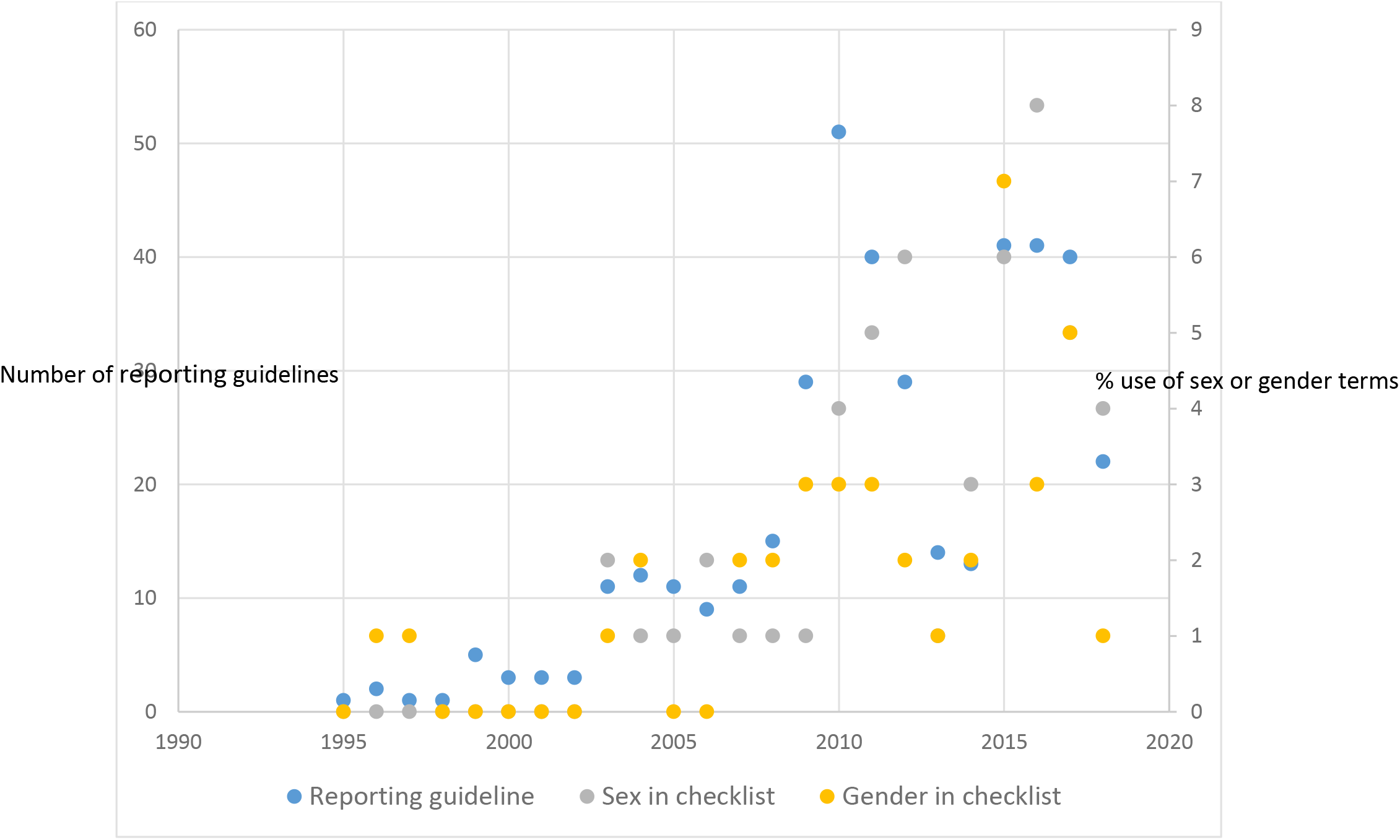
Publication trends in the use of sex and gender in the checklist

### Type of sex and gender information in the checklist

Of the reporting guidelines that mentioned “sex” and/or “gender” in their checklist, only two provided detailed information about “sex” (S1 Table, RG18, RG102) or “gender” (S1 Table, RG102). Two and four reporting guidelines specified the categories for “sex” and “gender”, respectively. The remaining vast majority just mentioned “sex” or “gender” terms in the list of demographic or baseline information.

## DISCUSSION

With the growing recognition of the importance of sex and gender considerations in research design and reporting, it is important to identify gaps in the integration of sex or gender considerations in reporting guidelines. We assessed the integration of sex and gender considerations in published reporting guidelines based on the use of sex and gender related words. At least one sex- and gender-related word was mentioned in slightly over half of the reporting guidelines assessed. In the checklist, the most important section of reporting guidelines, there was only 14% and 11% use of “sex” and “gender”, respectively; and 14% for “sex” and 18% for “gender” in the main text (statement). Only one reporting guideline correctly used “sex” and “gender” based on the criteria, and nonbinary was the least met criterion. Overall, “sex” and “gender” terms were increasingly used over time with an earlier use of “gender” than “sex”. Finally, only two reporting guidelines provided detailed information about “sex” or “gender” in their checklist. These findings lead us to the following observations.

We found that only a small number of reporting guidelines used “sex” and “gender” terms, particularly in their checklist. Studies that examined the use of sex- and gender-related words in published health research articles reported similar findings. In a review that described the considerations of “sex” and “gender” in 113 Cochrane reviews of interventions for preventing healthcare-associated infections (published between 2003 and 2016), 51% and 37% used “sex” and “gender”, respectively [21]. In another study that examined original investigations on diabetes, published in 2015 in the top ten general medicine and diabetes-specific journals based on impact factors, “sex” and “gender” were mentioned in the introduction section of 10% and in the methods sections of 30% [22]. More recently, “sex” was mentioned in 43% and “gender” in 41% of 87 studies (published between 1995 and 2017) included in a Cochrane review on the effectiveness of interventions for increasing the use of shared decision making by health professionals[19]. Thus, the proportion of the use of “sex” and “gender” terms in the reporting guidelines (published between 1995 and 2018) is far below that of original studies (14% versus 43% for “sex”; 18% versus 41% for “gender”) even though they were published during the same time period. Reporting guidelines are published as scientific articles that recommend the minimum elements required to adequately report different types of studies. Researchers, funders, and journal editors are among the end-users of reporting guidelines [2, 23], and there were significant moves on the importance of sex and gender considerations. For example, awareness articles and reports [24–27], requirements for grant applicants from top funders (Canadian Institutes of Health Research, the European Commission, and the US National Institutes of Health)[28–30], and specific instructions regarding sex and gender considerations by journal editors [31, 32]. Authors of reporting guidelines should play their part by integrating items regarding appropriate sex and gender considerations.

Only one reporting guideline met all three criteria of the correct use of “sex” and “gender” concepts. This is consistent with previous findings of studies that examined sex and gender considerations in health research, regardless of how “correct” or “appropriate” use was defined. In a previous review study, from which we adopted the definition of correct use (based on three criteria), no study met all three criteria [19]. Tannenbaum et al. found that only 35% of Canadian clinical practice guidelines (published between 2013 and 2015 for noncommunicable health conditions) that included “sex” and/or “gender” terms used them correctly according to the Sex and Gender Equity in Research Guidelines[5]. This result was expected, because only recently (in the 2010s) has a clear distinction between sex and gender, particularly in health research, reached the mainstream [25, 27, 29, 33–39]. Indeed, we found a modest increased use of “sex” and “gender” over time. The earlier use of “gender” in reporting guidelines was not surprising and might be used for “sex”, denoting the issues of inadequate and interchangeable use of these terms, particularly “gender”, as reported elsewhere [5, 19, 21, 22, 40]. These issues should be reflected in the future development of reporting guidelines or updates of existing guidelines.

Another aspect is the nature of the sex and gender information provided. In our study, detailed information was provided in only two checklists. As in similar studies [19, 21, 22], we were not able to assess the correct use of “sex” and “gender” for the vast majority of checklists (reporting guidelines) because of insufficient or inadequate information, even after consulting the explanation and elaboration documents of the reporting guidelines that used “sex” and “gender” terms in their checklist. Reporting guidelines are published as an article (statement) that most often includes a clear checklist of items to report. Pertinent to the publication of the Guidance for Developers of Health Research Reporting Guidelines (recommended steps for the development of reporting guidelines) [2], an increasing number of authors of reporting guidelines were publishing the companion explanation and elaboration documents. However, it is documented that very few authors actually consult those companion documents, hence, the suggestion to include more tailored and expanded details in the checklist of published reporting guidelines [17]. The proposed ‘change of paradigm’ would help authors of reporting guidelines to seize this opportunity to improve the integration of sex and gender by providing specific instructions on how to consider sex and/or gender within the different sections at the stage of writing the manuscripts. Alternatively, authors of reporting guidelines could harness the potential of information technology and work with the EQUATOR Network to integrate their reporting checklists into the platform that its partner company, Penelope, is developing to help authors identify relevant reporting guidelines. Indeed, the platform offers the flexibility to insert hyperlinks to detailed information[41].

Our study has some limitations. First, our systematic review was based on reporting guidelines in the Equator Network registry; thus, we cannot ensure that all published reporting guidelines of health research were included. However, we are confident that all the relevant reporting guidelines were included given the state-of-the-art processes of selection and curation by the EQUATOR Network Team. Second, we used the main document of reporting guidelines (statement) to assess the use of sex- and gender-related words and consulted only available explanation and elaboration documents for the reporting guidelines that used “sex” and “gender” terms in their checklist to assess correct use of these terms. This choice was justified by the fact that the checklist is the most important section of reporting guidelines, and authors rarely consult explanation and elaboration documents [17]. The comparison of electronic and manual counts of sex- and gender-related words showed that the electronic count, while not perfect, was the most accurate and is very unlikely to affect our assessment. We assessed evidence-based development of a sample of reporting guidelines due to practical constraints. However, by using a random sample, we are confident our results reflect the current quality of published reporting guidelines, with almost half of them not developed according to the recommended steps [2]. Finally, we assessed the presence of sex- and gender-related words but not how considerations of sex and gender were integrated into the guidelines. The appropriate level of sex and gender information in reporting guidelines remains to be determined. We can make the following recommendations:

- End users of current reporting guidelines should check the use of sex and gender terms against the standard definitions and make the necessary correction for their appropriate use while writing their manuscripts; they may also refer to SAGER [25].
- Journal editors should provide guidelines for transparent reporting of sex and gender [32], which is implemented by some journals such as JAMA.
- Authors of current reporting guidelines with obvious misuse of sex and gender concepts and related terms should address it by updating their statement [42].
- The EQUATOR Network should encourage developers of reporting guidelines (to consider sex and gender) and provide them with more operative information to consider sex and gender by updating their guidance[43].

## CONCLUSION

The integration of sex and gender considerations based on the use of sex- and gender-related words is very low in published reporting guidelines, particularly in their checklist. The initial step for authors of existing reporting guidelines remains to address the issues of inappropriate use of “sex” and “gender” concepts. Our findings will inform developers and users of these guidelines and may ultimately help reduce this gap for a better use of research knowledge and quality of evidence synthesis from health research.

## Supporting information caption

S1 Checklist. PRISMA checklist

S1 Figure. Publication trends in the mean number of occurrences of sex and gender in the checklist

S1 Table. List of included reporting guidelines

S2 Table. Comparison of electronic and manual identification of sex and gender related words in reporting guidelines

S3 Table. Concordance between electronic and manual identification of sex and gender related words in reporting guidelines

S4 Table. Distribution of the use of “sex” in various study types and sections of reporting guidelines

S5 Table. Distribution of the use of “gender” in various study types and sections of reporting guidelines

S1 Image. Example of manual count of sex- and gender-related words S1 Manual. Validity assessment for methods systematic review

S1 Database. Detailed internal validity assessment

