## Supplementary material for "Sex and Gender Considerations in Reporting Guidelines of Health Research: A Systematic Review": Gogovor_S1 Manual for Validity assessment for methods systematic review.docx

**Internal validity assessment for methods systematic reviews^[1]^**

1. Did the developers of the guideline represent more than one stakeholder group (e.g. researchers, funders, publishers)?

*Note: Look for anything about stakeholder consultations in the text of the paper.*

*An answer of yes = they explicitly reported more than one stakeholder group; if a specific group is named (e.g., the CONSORT Group), verify its composition on the internet.*

*An answer of no = they were explicit and reported that they did not represent other stakeholders. Otherwise answer = unclear.*

2. Did the developers report gathering any data for the creation of the guideline (e.g., carry out a literature review, collect anecdotal data)?

*Note: methods of data gathering must be clearly stated. Citations next to checklist items do not count as data gathering. Look for information in the text only.*

*An answer of yes = they were explicit mention of any type of data gathering*

*An answer of no = they were explicit and reported that they did not gather any data. Otherwise answer = unclear.*

3. Did the developers report the use of a consensus process (e.g., Delphi, RAND/UCLA Appropriateness Method, nominal group technique, consensus, meeting, development conference, roundtable, panel, in-person discussion)?

*Note:*

*An answer of yes = they were explicit and reported any of the consensus activities listed above*

*An answer of no = they were explicit and reported that they did not conduct any consensus activities. Otherwise answer = unclear.*

1. Adapted from Cukier S, Helal L, Rice DB, Pupkaite J, Ahmadzai N, Wilson M, et al. Checklists to detect potential predatory biomedical journals: a systematic review. BMC Med. 2020; 18(1):104.

   **Goal**: To assess evidence-based development of the reporting guideline

   **Scoring**:	“yes” (i.e. low risk of bias), “no” (i.e. high risk of bias) or “unclear” (i.e. unclear risk of bias).

   **Overall**:	high internal validity (low risk of bias): ≥ 2 “yes”

   low internal validity (high risk of bias): < 2 “yes”
