## Supplementary material for "Sex and Gender Considerations in Reporting Guidelines of Health Research: A Systematic Review": Gogovor_Supplement documents_Sex and gender considerations in reporting guidelines.docx

**Gogovor et al. *PLOS Medicine***

mean number of use of the word

| year | Gender - check | Sex - check |
| --- | --- | --- |
| 1995 | 0 | 0 |
| 1996 | #N/A | 0 |
| 1997 | #N/A | 0 |
| 1998 | 0 | 0 |
| 1999 | 0 | 0 |
| 2000 | 0 | 0 |
| 2001 | 0 | 0 |
| 2002 | 0 | 0 |
| 2003 | 0.0909091 | 0.0909091 |
| 2004 | 0.166667 | 0.0833333 |
| 2005 | 0 | 0.0909091 |
| 2006 | 0 | 0.555556 |
| 2007 | 0.181818 | 0.0909091 |
| 2008 | 0.133333 | 0.0666667 |
| 2009 | 0.103448 | 0.0344828 |
| 2010 | 0.0980392 | 0.0980392 |
| 2011 | 0.075 | 0.125 |
| 2012 | 0.0689655 | 0.206897 |
| 2013 | 0.0714286 | 0.142857 |
| 2014 | 0.153846 | 0.384615 |
| 2015 | 0.243902 | 0.170732 |
| 2016 | 0.317073 | 0.585366 |
| 2017 | 0.35 | 0.375 |
| 2018 | 0.0454545 | 0.545454 |

S1 Figure. Publication trends in the mean number of occurrences of sex and gender in the checklist

**Supplement Table 1. List of included reporting guidelines**

| **No** | **Reference** | **Acronym** | **Use of “sex” term in checklist** | **Use of “gender” term in checklist** |
| --- | --- | --- | --- | --- |
| RG1 | af Wåhlberg AE. A reporting guide for studies on individual differences in traffic safety. Journal of Safety Research. 2010;41(4):381-3. |  |  |  |
| RG2 | Agarwal S, LeFevre AE, Lee J, L'Engle K, Mehl G, Sinha C, et al. Guidelines for reporting of health interventions using mobile phones: mobile health (mHealth) evidence reporting and assessment (mERA) checklist. BMJ (Clinical research ed). 2016;352:i1174. | mERA |  |  |
| RG3 | Agha RA, Borrelli MR, Farwana R, Koshy K, Fowler AJ, Orgill DP. The PROCESS 2018 statement: Updating Consensus Preferred Reporting Of CasE Series in Surgery (PROCESS) guidelines. International journal of surgery (London, England). 2018;60:279-82. | PROCESS |  |  |
| RG4 | Agha RA, Borrelli MR, Farwana R, Koshy K, Fowler AJ, Orgill DP. The SCARE 2018 statement: Updating consensus Surgical CAse REport (SCARE) guidelines. International journal of surgery (London, England). 2018;60:132-6. | SCARE |  |  |
| RG5 | Agha RA, Borrelli MR, Vella-Baldacchino M, Thavayogan R, Orgill DP. The STROCSS statement: Strengthening the Reporting of Cohort Studies in Surgery. International journal of surgery (London, England). 2017;46:198-202. | STROCSS |  |  |
| RG6 | Ahmed M, Solbiati L, Brace CL, Breen DJ, Callstrom MR, Charboneau JW, et al. Image-guided tumor ablation: standardization of terminology and reporting criteria--a 10-year update. Radiology. 2014;273(1):241-60. |  |  |  |
| RG7 | Akard LP, Wang YL. Translating trial-based molecular monitoring into clinical practice: importance of international standards and practical considerations for community practitioners. Clinical lymphoma, myeloma & leukemia. 2011;11(5):385-95. |  |  |  |
| RG8 | Albrecht L, Archibald M, Arseneau D, Scott SD. WIDER recommendations for reporting of behaviour change interventions. On behalf of the Workgroup for Intervention Development and Evaluation Research. Implementation science : IS. 2013;8(52). | WIDER |  |  |
| RG9 | Aletaha D, Landewe R, Karonitsch T, Bathon J, Boers M, Bombardier C, et al. Reporting disease activity in clinical trials of patients with rheumatoid arthritis: EULAR/ACR collaborative recommendations. Annals of the rheumatic diseases. 2008;67(10):1360-4. |  |  |  |
| RG10 | Andrews NA, Latrémolière A, Basbaum AI, Mogil JS, Porreca F, Rice ASC, et al. Ensuring transparency and minimization of methodologic bias in preclinical pain research: PPRECISE considerations. Pain. 2016;157(4):901-9. |  |  |  |
| RG11 | Armstrong D, Barkun A, Bridges R, Carter R, de Gara C, Dube C, et al. Canadian Association of Gastroenterology consensus guidelines on safety and quality indicators in endoscopy. Canadian journal of gastroenterology = Journal canadien de gastroenterologie. 2012;26(1):17-31. |  |  |  |
| RG12 | Aronson JK. Anecdotes as evidence. BMJ (Clinical research ed). 2003;326(7403):1346-. |  |  |  |
| RG13 | Arroz M, Came N, Lin P, Chen W, Yuan C, Lagoo A, et al. Consensus guidelines on plasma cell myeloma minimal residual disease analysis and reporting. Cytometry Part B, Clinical cytometry. 2016;90(1):31-9. |  |  |  |
| RG14 | Atkinson KM, Koenka AC, Sanchez CE, Moshontz H, Cooper H. Reporting standards for literature searches and report inclusion criteria: making research syntheses more transparent and easy to replicate. Research synthesis methods. 2015;6(1):87-95. |  |  |  |
| RG15 | Baker TB, Gustafson DH, Shaw B, Hawkins R, Pingree S, Roberts L, et al. Relevance of CONSORT reporting criteria for research on eHealth interventions. Patient education and counseling. 2010;81 Suppl:S77-86. |  |  |  |
| RG16 | Baldwin SA, Larson MJ. An introduction to using Bayesian linear regression with clinical data. Behaviour research and therapy. 2017;98:58-75. |  |  |  |
| RG17 | Banovac F, Buckley DC, Kuo WT, Lough DM, Martin LG, Millward SF, et al. Reporting standards for endovascular treatment of pulmonary embolism. Journal of vascular and interventional radiology : JVIR. 2010;21(1):44-53. |  |  |  |
| RG18 | Battisti WP, Wager E, Baltzer L, Bridges D, Cairns A, Carswell CI, et al. Good Publication Practice for Communicating Company-Sponsored Medical Research: GPP3. Annals of internal medicine. 2015;163(6):461-4. |  |  |  |
| RG19 | Beller EM, Glasziou PP, Altman DG, Hopewell S, Bastian H, Chalmers I, et al. PRISMA for Abstracts: reporting systematic reviews in journal and conference abstracts. PLoS medicine. 2013;10(4):e1001419. |  |  |  |
| RG20 | Benchimol EI, Manuel DG, To T, Griffiths AM, Rabeneck L, Guttmann A. Development and use of reporting guidelines for assessing the quality of validation studies of health administrative data. Journal of clinical epidemiology. 2011;64(8):821-9. |  |  |  |
| RG21 | Benchimol EI, Smeeth L, Guttmann A, Harron K, Moher D, Petersen I, et al. The REporting of studies Conducted using Observational Routinely-collected health Data (RECORD) statement. PLoS medicine. 2015;12(10):e1001885. | RECORD |  |  |
| RG22 | Bennett DA, Brayne C, Feigin VL, Barker-Collo S, Brainin M, Davis D, et al. Development of the standards of reporting of neurological disorders (STROND) checklist: a guideline for the reporting of incidence and prevalence studies in neuroepidemiology. European journal of epidemiology. 2015;30(7):569-76. | STROND |  |  |
| RG23 | Berger AM, Wielgus KK, Young-McCaughan S, Fischer P, Farr L, Lee KA. Methodological challenges when using actigraphy in research. Journal of pain and symptom management. 2008;36(2):191-9. |  |  |  |
| RG24 | Berger ML, Mamdani M, Atkins D, Johnson ML. Good research practices for comparative effectiveness research: defining, reporting and interpreting nonrandomized studies of treatment effects using secondary data sources: the ISPOR Good Research Practices for Retrospective Database Analysis Task Force Report--Part I. Value in health : the journal of the International Society for Pharmacoeconomics and Outcomes Research. 2009;12(8):1044-52. |  |  |  |
| RG25 | Biering-Sorensen F, Charlifue S, Devivo MJ, Grinnon ST, Kleitman N, Lu Y, et al. Using the Spinal Cord Injury Common Data Elements. Topics in spinal cord injury rehabilitation. 2012;18(1):23-7. |  |  |  |
| RG26 | Biesecker LG. An introduction to standardized clinical nomenclature for dysmorphic features: the Elements of Morphology project. BMC medicine. 2010;8:56. |  |  |  |
| RG27 | Bishop C, Paul G, Thewlis D. Recommendations for the reporting of foot and ankle models. Journal of biomechanics. 2012;45(13):2185-94. |  |  |  |
| RG28 | Black CM, Thorpe K, Venrbux A, Kim HS, Millward SF, Clark TW, et al. Research reporting standards for endovascular treatment of pelvic venous insufficiency. Journal of vascular and interventional radiology : JVIR. 2010;21(6):796-803. |  |  |  |
| RG29 | Blignault I, Ritchie J. Revealing the wood and the trees: reporting qualitative research. Health promotion journal of Australia : official journal of Australian Association of Health Promotion Professionals. 2009;20(2):140-5. |  |  |  |
| RG30 | Bodemer N, Muller SM, Okan Y, Garcia-Retamero R, Neumeyer-Gromen A. Do the media provide transparent health information? A cross-cultural comparison of public information about the HPV vaccine. Vaccine. 2012;30(25):3747-56. |  |  |  |
| RG31 | Boers M. Graphics and statistics for cardiology: designing effective tables for presentation and publication. Heart (British Cardiac Society). 2018;104(3):192-200. |  |  |  |
| RG32 | Bohensky MA, Jolley D, Sundararajan V, Evans S, Ibrahim J, Brand C. Development and validation of reporting guidelines for studies involving data linkage. Australian and New Zealand journal of public health. 2011;35(5):486-9. |  |  |  |
| RG33 | Boller M, Fletcher DJ, Brainard BM, Haskins S, Hopper K, Nadkarni VM, et al. Utstein-style guidelines on uniform reporting of in-hospital cardiopulmonary resuscitation in dogs and cats. A RECOVER statement. Journal of veterinary emergency and critical care (San Antonio, Tex : 2001). 2016;26(1):11-34. |  |  |  |
| RG34 | Bookman EB, Langehorne AA, Eckfeldt JH, Glass KC, Jarvik GP, Klag M, et al. Reporting genetic results in research studies: summary and recommendations of an NHLBI working group. American journal of medical genetics Part A. 2006;140(10):1033-40. |  |  |  |
| RG35 | Booth A. "Brimful of STARLITE": toward standards for reporting literature searches. Journal of the Medical Library Association : JMLA. 2006;94(4):421-9, e205. |  |  |  |
| RG36 | Bordeianu CD, Ticu CE. A new manner of reporting pressure results after glaucoma surgery. Clinical ophthalmology (Auckland, NZ). 2012;6:23-31. |  |  |  |
| RG37 | Borek AJ, Abraham C, Smith JR, Greaves CJ, Tarrant M. A checklist to improve reporting of group-based behaviour-change interventions. BMC public health. 2015;15:963. |  |  |  |
| RG38 | Bossuyt PM, Reitsma JB, Bruns DE, Gatsonis CA, Glasziou PP, Irwig L, et al. STARD 2015: An Updated List of Essential Items for Reporting Diagnostic Accuracy Studies. Radiology. 2015;277(3):826-32. | STARD 2015 |  |  |
| RG39 | Bougioukas KI, Bouras E, Apostolidou-Kiouti F, Kokkali S, Arvanitidou M, Haidich AB. Reporting guidelines on how to write a complete and transparent abstract for overviews of systematic reviews of health care interventions. Journal of clinical epidemiology. 2018. | PRIO for abstracts |  |  |
| RG40 | Bougioukas KI, Liakos A, Tsapas A, Ntzani E, Haidich AB. Preferred reporting items for overviews of systematic reviews including harms checklist: a pilot tool to be used for balanced reporting of benefits and harms. Journal of clinical epidemiology. 2018;93:9-24. | PRIO-harms |  |  |
| RG41 | Boursier J, de Ledinghen V, Poynard T, Guechot J, Carrat F, Leroy V, et al. An extension of STARD statements for reporting diagnostic accuracy studies on liver fibrosis tests: the Liver-FibroSTARD standards. Journal of hepatology. 2015;62(4):807-15. | Liver-FibroSTARD |  |  |
| RG42 | Bousquet PJ, Brozek J, Bachert C, Bieber T, Bonini S, Burney P, et al. The CONSORT statement checklist in allergen-specific immunotherapy: a GA2LEN paper. Allergy. 2009;64(12):1737-45. |  |  |  |
| RG43 | Boutron I, Altman DG, Moher D, Schulz KF, Ravaud P. CONSORT Statement for Randomized Trials of Nonpharmacologic Treatments: A 2017 Update and a CONSORT Extension for Nonpharmacologic Trial Abstracts. Annals of internal medicine. 2017;167(1):40-7. | CONSORT Non-pharmacologic treatment intervention |  |  |
| RG44 | Bouxsein ML, Boyd SK, Christiansen BA, Guldberg RE, Jepsen KJ, Muller R. Guidelines for assessment of bone microstructure in rodents using micro-computed tomography. Journal of bone and mineral research : the official journal of the American Society for Bone and Mineral Research. 2010;25(7):1468-86. |  |  |  |
| RG45 | Bradt DA, Aitken P. Disaster medicine reporting: the need for new guidelines and the CONFIDE statement. Emergency medicine Australasia : EMA. 2010;22(6):483-7. | CONFIDE |  |  |
| RG46 | Bramhall M, Florez-Vargas O, Stevens R, Brass A, Cruickshank S. Quality of methods reporting in animal models of colitis. Inflammatory bowel diseases. 2015;21(6):1248-59. |  |  |  |
| RG47 | Bravo E, Calzolari A, De Castro P, Mabile L, Napolitani F, Rossi AM, et al. Developing a guideline to standardize the citation of bioresources in journal articles (CoBRA). BMC medicine. 2015;13:33. | CoBRA |  |  |
| RG48 | Brethauer SA, Kim J, el Chaar M, Papasavas P, Eisenberg D, Rogers A, et al. Standardized outcomes reporting in metabolic and bariatric surgery. Surgery for obesity and related diseases : official journal of the American Society for Bariatric Surgery. 2015;11(3):489-506. |  |  |  |
| RG49 | Breuer E, Lee L, De Silva M, Lund C. Using theory of change to design and evaluate public health interventions: a systematic review. Implementation science : IS. 2016;11:63. |  |  |  |
| RG50 | Brookhart MA, Rassen JA, Schneeweiss S. Instrumental variable methods in comparative safety and effectiveness research. Pharmacoepidemiology and drug safety. 2010;19(6):537-54. |  |  |  |
| RG51 | Brouwers MC, Kerkvliet K, Spithoff K. The AGREE Reporting Checklist: a tool to improve reporting of clinical practice guidelines. BMJ (Clinical research ed). 2016;352:i1152. | AGREE |  |  |
| RG52 | Brown DB, Gould JE, Gervais DA, Goldberg SN, Murthy R, Millward SF, et al. Transcatheter therapy for hepatic malignancy: standardization of terminology and reporting criteria. Journal of vascular and interventional radiology : JVIR. 2009;20(7 Suppl):S425-34. |  |  |  |
| RG53 | Brown P, Brunnhuber K, Chalkidou K, Chalmers I, Clarke M, Fenton M, et al. How to formulate research recommendations. BMJ (Clinical research ed). 2006;333(7572):804-6. |  |  |  |
| RG54 | Bruck K, Jager KJ, Dounousi E, Kainz A, Nitsch D, Arnlov J, et al. Methodology used in studies reporting chronic kidney disease prevalence: a systematic literature review. Nephrology, dialysis, transplantation : official publication of the European Dialysis and Transplant Association - European Renal Association. 2015;30 Suppl 4:iv6-16. |  |  |  |
| RG55 | Burns KE, Duffett M, Kho ME, Meade MO, Adhikari NK, Sinuff T, et al. A guide for the design and conduct of self-administered surveys of clinicians. CMAJ : Canadian Medical Association journal = journal de l'Association medicale canadienne. 2008;179(3):245-52. |  |  |  |
| RG56 | Burton A, Altman DG. Missing covariate data within cancer prognostic studies: a review of current reporting and proposed guidelines. British journal of cancer. 2004;91(1):4-8. |  |  |  |
| RG57 | Callstrom MR, York JD, Gaba RC, Gemmete JJ, Gervais DA, Millward SF, et al. Research reporting standards for image-guided ablation of bone and soft tissue tumors. Journal of vascular and interventional radiology : JVIR. 2009;20(12):1527-40. |  |  |  |
| RG58 | Calvert M, Blazeby J, Altman DG, Revicki DA, Moher D, Brundage MD. Reporting of patient-reported outcomes in randomized trials: the CONSORT PRO extension. Jama. 2013;309(8):814-22. | CONSORT-PRO |  |  |
| RG59 | Calvert M, Kyte D, Mercieca-Bebber R, Slade A, Chan AW, King MT, et al. Guidelines for Inclusion of Patient-Reported Outcomes in Clinical Trial Protocols: The SPIRIT-PRO Extension. Jama. 2018;319(5):483-94. | SPIRIT-PRO |  |  |
| RG60 | Cambon L, Minary L, Ridde V, Alla F. A tool to analyze the transferability of health promotion interventions. BMC public health. 2013;13:1184. | ASTAIRE |  |  |
| RG61 | Campana LG, Clover AJ, Valpione S, Quaglino P, Gehl J, Kunte C, et al. Recommendations for improving the quality of reporting clinical electrochemotherapy studies based on qualitative systematic review. Radiology and oncology. 2016;50(1):1-13. |  |  |  |
| RG62 | Campbell M, Katikireddi SV, Hoffmann T, Armstrong R, Waters E, Craig P. TIDieR-PHP: a reporting guideline for population health and policy interventions. BMJ (Clinical research ed). 2018;361:k1079. | TIDieR-PHP |  |  |
| RG63 | Campbell MK, Piaggio G, Elbourne DR, Altman DG. Consort 2010 statement: extension to cluster randomised trials. BMJ (Clinical research ed). 2012;345:e5661. | CONSORT Cluster |  |  |
| RG64 | Carracedo A, Butler JM, Gusmao L, Parson W, Roewer L, Schneider PM. Publication of population data for forensic purposes. Forensic science international Genetics. 2010;4(3):145-7. |  |  |  |
| RG65 | Castren M, Bohm K, Kvam AM, Bovim E, Christensen EF, Steen-Hansen JE, et al. Reporting of data from out-of-hospital cardiac arrest has to involve emergency medical dispatching--taking the recommendations on reporting OHCA the Utstein style a step further. Resuscitation. 2011;82(12):1496-500. |  |  |  |
| RG66 | Castren M, Karlsten R, Lippert F, Christensen EF, Bovim E, Kvam AM, et al. Recommended guidelines for reporting on emergency medical dispatch when conducting research in emergency medicine: the Utstein style. Resuscitation. 2008;79(2):193-7. |  |  |  |
| RG67 | Chan AW, Tetzlaff JM, Altman DG, Laupacis A, Gotzsche PC, Krleza-Jeric K, et al. SPIRIT 2013 statement: defining standard protocol items for clinical trials. Annals of internal medicine. 2013;158(3):200-7. | SPIRIT 2013 |  |  |
| RG68 | Chang A, Gibson IW, Cohen AH, Weening JW, Jennette JC, Fogo AB. A position paper on standardizing the nonneoplastic kidney biopsy report. Human pathology. 2012;43(8):1192-6. |  |  |  |
| RG69 | Chang S, Vogelbaum M, Lang FF, Haines S, Kunwar S, Chiocca EA, et al. GNOSIS: guidelines for neuro-oncology: standards for investigational studies--reporting of surgically based therapeutic clinical trials. Journal of neuro-oncology. 2007;82(2):211-20. |  |  |  |
| RG70 | Chang SM, Reynolds SL, Butowski N, Lamborn KR, Buckner JC, Kaplan RS, et al. GNOSIS: guidelines for neuro-oncology: standards for investigational studies-reporting of phase 1 and phase 2 clinical trials. Neuro-oncology. 2005;7(4):425-34. |  |  |  |
| RG71 | Chen Y, Yang K, Marusic A, Qaseem A, Meerpohl JJ, Flottorp S, et al. A Reporting Tool for Practice Guidelines in Health Care: The RIGHT Statement. Annals of internal medicine. 2017;166(2):128-32. | RIGHT |  |  |
| RG72 | Cheng A, Kessler D, Mackinnon R, Chang TP, Nadkarni VM, Hunt EA, et al. Reporting Guidelines for Health Care Simulation Research: Extensions to the CONSORT and STROBE Statements. Simulation in healthcare : journal of the Society for Simulation in Healthcare. 2016;11(4):238-48. |  |  |  |
| RG73 | Cheng CW, Wu TX, Shang HC, Li YP, Altman DG, Moher D, et al. CONSORT Extension for Chinese Herbal Medicine Formulas 2017: Recommendations, Explanation, and Elaboration. Annals of internal medicine. 2017;167(2):112-21. | CONSORT-CHM |  |  |
| RG74 | Cheson BD, Bennett JM, Kopecky KJ, Buchner T, Willman CL, Estey EH, et al. Revised recommendations of the International Working Group for Diagnosis, Standardization of Response Criteria, Treatment Outcomes, and Reporting Standards for Therapeutic Trials in Acute Myeloid Leukemia. Journal of clinical oncology : official journal of the American Society of Clinical Oncology. 2003;21(24):4642-9. |  |  |  |
| RG75 | Chipperfield L, Citrome L, Clark J, David FS, Enck R, Evangelista M, et al. Authors' Submission Toolkit: a practical guide to getting your research published. Current medical research and opinion. 2010;26(8):1967-82. |  |  |  |
| RG76 | Choi J, Choi T-Y, Jun JH, Lee JA, Lee MS. Preferred Reporting Items for the Development of Evidence-based Clinical Practice Guidelines in Traditional Medicine (PRIDE-CPG-TM): Explanation and elaboration. European Journal of Integrative Medicine. 2016;8(6):905-15. | PRIDE-CPG-TM |  |  |
| RG77 | Christopher MM, Young KM. Writing for Publication in Veterinary Medicine. A Practical Guide for Researchers and Clinicians: Wiley-Blackwell; 2011. Available from: http://eu.wiley.com/WileyCDA/Section/id-612222.html. |  |  |  |
| RG78 | Clark JP. How to Peer Review a Qualitative Manuscript. In: Godlee F, Jefferson T, editors. Peer Review in Health Sciences. 2. London: BMJ Books; 2003. p. 219-35. |  |  |  |
| RG79 | Clark TW, Millward SF, Gervais DA, Goldberg SN, Grassi CJ, Kinney TB, et al. Reporting standards for percutaneous thermal ablation of renal cell carcinoma. Journal of vascular and interventional radiology : JVIR. 2009;20(7 Suppl):S409-16. |  |  |  |
| RG80 | Coast J, Al-Janabi H, Sutton EJ, Horrocks SA, Vosper AJ, Swancutt DR, et al. Using qualitative methods for attribute development for discrete choice experiments: issues and recommendations. Health economics. 2012;21(6):730-41. |  |  |  |
| RG81 | Cohen JF, Korevaar DA, Gatsonis CA, Glasziou PP, Hooft L, Moher D, et al. STARD for Abstracts: essential items for reporting diagnostic accuracy studies in journal or conference abstracts. BMJ (Clinical research ed). 2017;358:j3751. | STARD for Abstracts |  |  |
| RG82 | Colbert AP, Spaulding K, Larsen A, Ahn AC, Cutro JA. Electrodermal activity at acupoints: literature review and recommendations for reporting clinical trials. Journal of acupuncture and meridian studies. 2011;4(1):5-13. |  |  |  |
| RG83 | Cole TJ. Setting number of decimal places for reporting risk ratios: rule of four. BMJ (Clinical research ed). 2015;350:h1845. |  |  |  |
| RG84 | Cole TJ. Too many digits: the presentation of numerical data. Archives of disease in childhood. 2015;100(7):608-9. |  |  |  |
| RG85 | Comenzo RL, Reece D, Palladini G, Seldin D, Sanchorawala V, Landau H, et al. Consensus guidelines for the conduct and reporting of clinical trials in systemic light-chain amyloidosis. Leukemia. 2012;26(11):2317-25. |  |  |  |
| RG86 | Conn VS, Groves PS. Protecting the power of interventions through proper reporting. Nursing outlook. 2011;59(6):318-25. |  |  |  |
| RG87 | Cook JA, Hislop J, Altman DG, Fayers P, Briggs AH, Ramsay CR, et al. Specifying the target difference in the primary outcome for a randomised controlled trial: guidance for researchers. Trials. 2015;16:12. |  |  |  |
| RG88 | Cook JL, Evans R, Conzemius MG, Lascelles BD, McIlwraith CW, Pozzi A, et al. Proposed definitions and criteria for reporting time frame, outcome, and complications for clinical orthopedic studies in veterinary medicine. Veterinary surgery : VS. 2010;39(8):905-8. |  |  |  |
| RG89 | Coroneos CJ, Ignacy TA, Thoma A. Designing and reporting case series in plastic surgery. Plastic and reconstructive surgery. 2011;128(4):361e-8e. |  |  |  |
| RG90 | Crawford JR, Garthwaite PH, Porter S. Point and interval estimates of effect sizes for the case-controls design in neuropsychology: rationale, methods, implementations, and proposed reporting standards. Cognitive neuropsychology. 2010;27(3):245-60. |  |  |  |
| RG91 | Creinin MD, Chen MJ. Medical abortion reporting of efficacy: the MARE guidelines. Contraception. 2016;94(2):97-103. | MARE-C & MARE-S |  |  |
| RG92 | Cummins RO, Chamberlain D, Hazinski MF, Nadkarni V, Kloeck W, Kramer E, et al. Recommended guidelines for reviewing, reporting, and conducting research on in-hospital resuscitation: the in-hospital 'Utstein style'. A statement for healthcare professionals from the American Heart Association, the European Resuscitation Council, the Heart and Stroke Foundation of Canada, the Australian Resuscitation Council, and the Resuscitation Councils of Southern Africa. Resuscitation. 1997;34(2):151-83. |  |  |  |
| RG93 | Currow DC, Tieman JJ, Greene A, Zafar SY, Wheeler JL, Abernethy AP. Refining a checklist for reporting patient populations and service characteristics in hospice and palliative care research. Journal of pain and symptom management. 2012;43(5):902-10. |  |  |  |
| RG94 | Czigany Z, Iwasaki J, Yagi S, Nagai K, Szijarto A, Uemoto S, et al. Improving Research Practice in Rat Orthotopic and Partial Orthotopic Liver Transplantation: A Review, Recommendation, and Publication Guide. European surgical research Europaische chirurgische Forschung Recherches chirurgicales europeennes. 2015;55(1-2):119-38. |  |  |  |
| RG95 | Darcourt J, Booij J, Tatsch K, Varrone A, Vander Borght T, Kapucu OL, et al. EANM procedure guidelines for brain neurotransmission SPECT using (123)I-labelled dopamine transporter ligands, version 2. European journal of nuclear medicine and molecular imaging. 2010;37(2):443-50. |  |  |  |
| RG96 | Davidson KW, Goldstein M, Kaplan RM, Kaufmann PG, Knatterud GL, Orleans CT, et al. Evidence-based behavioral medicine: what is it and how do we achieve it? Annals of behavioral medicine : a publication of the Society of Behavioral Medicine. 2003;26(3):161-71. |  |  |  |
| RG97 | Davis JC, Robertson MC, Comans T, Scuffham PA. Guidelines for conducting and reporting economic evaluation of fall prevention strategies. Osteoporosis international: a journal established as result of cooperation between the European Foundation for Osteoporosis and the National Osteoporosis Foundation of the USA. 2011;22(9):2449-59. |  |  |  |
| RG98 | Davis MF, Rankin SC, Schurer JM, Cole S, Conti L, Rabinowitz P, et al. Checklist for One Health Epidemiological Reporting of Evidence (COHERE). One Health. 2017;4:14-21. | COHERE |  |  |
| RG99 | De Geest S, Zullig LL, Dunbar-Jacob J, Helmy R, Hughes DA, Wilson IB, et al. ESPACOMP Medication Adherence Reporting Guideline (EMERGE). Annals of internal medicine. 2018;169(1):30-5. | EMERGE |  |  |
| RG100 | de Jager DJ, de Mutsert R, Jager KJ, Zoccali C, Dekker FW. Reporting of interaction. Nephron Clinical practice. 2011;119(2):c158-61. |  |  |  |
| RG101 | de Keizer NF, Talmon J, Ammenwerth E, Brender J, Rigby M, Nykanen P. Systematic prioritization of the STARE-HI reporting items. An application to short conference papers on health informatics evaluation. Methods of information in medicine. 2012;51(2):104-11. |  |  |  |
| RG102 | de Vries RBM, Hooijmans CR, Langendam MW, van Luijk J, Leenaars M, Ritskes-Hoitinga M, et al. A protocol format for the preparation, registration and publication of systematic reviews of animal intervention studies. 2015;2(1):e00007. |  |  |  |
| RG103 | Dean ME, Coulter MK, Fisher P, Jobst KA, Walach H. Reporting data on homeopathic treatments (RedHot): a supplement to CONSORT. Journal of alternative and complementary medicine (New York, NY). 2007;13(1):19-23. |  |  |  |
| RG104 | Dechartres A, Boutron I, Roy C, Ravaud P. Inadequate planning and reporting of adjudication committees in clinical trials: recommendation proposal. Journal of clinical epidemiology. 2009;62(7):695-702. |  |  |  |
| RG105 | Des Jarlais DC, Lyles C, Crepaz N. Improving the reporting quality of nonrandomized evaluations of behavioral and public health interventions: the TREND statement. American journal of public health. 2004;94(3):361-6. | TREND |  |  |
| RG106 | DeVivo MJ, Biering-Sorensen F, New P, Chen Y. Standardization of data analysis and reporting of results from the International Spinal Cord Injury Core Data Set. Spinal cord. 2011;49(5):596-9. |  |  |  |
| RG107 | Dick WF, Baskett PJ. Recommendations for uniform reporting of data following major trauma--the Utstein style. A report of a working party of the International Trauma Anaesthesia and Critical Care Society (ITACCS). Resuscitation. 1999;42(2):81-100. |  |  |  |
| RG108 | Dionne CE, Dunn KM, Croft PR, Nachemson AL, Buchbinder R, Walker BF, et al. A consensus approach toward the standardization of back pain definitions for use in prevalence studies. Spine. 2008;33(1):95-103. | DOLBaPP |  |  |
| RG109 | Dixon WG, Carmona L, Finckh A, Hetland ML, Kvien TK, Landewe R, et al. EULAR points to consider when establishing, analysing and reporting safety data of biologics registers in rheumatology. Annals of the rheumatic diseases. 2010;69(9):1596-602. |  |  |  |
| RG110 | Docherty M, Smith R. The case for structuring the discussion of scientific papers. BMJ (Clinical research ed). 1999;318(7193):1224-5. |  |  |  |
| RG111 | Dohner H, Estey EH, Amadori S, Appelbaum FR, Buchner T, Burnett AK, et al. Diagnosis and management of acute myeloid leukemia in adults: recommendations from an international expert panel, on behalf of the European LeukemiaNet. Blood. 2010;115(3):453-74. |  |  |  |
| RG112 | Donahue SP, Arnold RW, Ruben JB. Preschool vision screening: what should we be detecting and how should we report it? Uniform guidelines for reporting results of preschool vision screening studies. Journal of AAPOS : the official publication of the American Association for Pediatric Ophthalmology and Strabismus. 2003;7(5):314-6. |  |  |  |
| RG113 | Dougados M, Simon P, Braun J, Burgos-Vargas R, Maksymowych WP, Sieper J, et al. ASAS recommendations for collecting, analysing and reporting NSAID intake in clinical trials/epidemiological studies in axial spondyloarthritis. Annals of the rheumatic diseases. 2011;70(2):249-51. |  |  |  |
| RG114 | Drummond M, Houwing N, Slothuus U, Giangrande P. Making economic evaluations more helpful for treatment choices in haemophilia. Haemophilia : the official journal of the World Federation of Hemophilia. 2017;23(2):e58-e66. |  |  |  |
| RG115 | Drummond M, Manca A, Sculpher M. Increasing the generalizability of economic evaluations: recommendations for the design, analysis, and reporting of studies. International journal of technology assessment in health care. 2005;21(2):165-71. |  |  |  |
| RG116 | Duffis EJ, Gandhi CD, Prestigiacomo CJ, Abruzzo T, Albuquerque F, Bulsara KR, et al. Head, neck, and brain tumor embolization guidelines. Journal of neurointerventional surgery. 2012;4(4):251-5. |  |  |  |
| RG117 | Dykstra K, Mehrotra N, Tornoe CW, Kastrissios H, Patel B, Al-Huniti N, et al. Reporting guidelines for population pharmacokinetic analyses. Journal of pharmacokinetics and pharmacodynamics. 2015;42(3):301-14. |  |  |  |
| RG118 | Eldridge SM, Chan CL, Campbell MJ, Bond CM, Hopewell S, Thabane L, et al. CONSORT 2010 statement: extension to randomised pilot and feasibility trials. BMJ (Clinical research ed). 2016;355:i5239. |  |  |  |
| RG119 | Elliott R, Fischer CT, Rennie DL. Evolving guidelines for publication of qualitative research studies in psychology and related fields. The British journal of clinical psychology. 1999;38 (Pt 3):215-29. |  |  |  |
| RG120 | Eysenbach G. Improving the quality of Web surveys: the Checklist for Reporting Results of Internet E-Surveys (CHERRIES). Journal of medical Internet research. 2004;6(3):e34. | CHERRIES |  |  |
| RG121 | Eysenbach G. CONSORT-EHEALTH: improving and standardizing evaluation reports of Web-based and mobile health interventions. Journal of medical Internet research. 2011;13(4):e126. | CONSORT-EHEALTH |  |  |
| RG122 | Feneck R, Kneeshaw J, Fox K, Bettex D, Erb J, Flaschkampf F, et al. Recommendations for reporting perioperative transoesophageal echo studies. European journal of echocardiography : the journal of the Working Group on Echocardiography of the European Society of Cardiology. 2010;11(5):387-93. |  |  |  |
| RG123 | Fernandes RM, van der Lee JH, Offringa M. A systematic review of the reporting of Data Monitoring Committees' roles, interim analysis and early termination in pediatric clinical trials. BMC pediatrics. 2009;9:77-. |  |  |  |
| RG124 | Field N, Cohen T, Struelens MJ, Palm D, Cookson B, Glynn JR, et al. Strengthening the Reporting of Molecular Epidemiology for Infectious Diseases (STROME-ID): an extension of the STROBE statement. The Lancet Infectious diseases. 2014;14(4):341-52. | STROME-ID |  |  |
| RG125 | Fillinger MF, Greenberg RK, McKinsey JF, Chaikof EL. Reporting standards for thoracic endovascular aortic repair (TEVAR). Journal of vascular surgery. 2010;52(4):1022-33, 33.e15. | TEVAR |  |  |
| RG126 | First draft of the STROBE checklist of items to be included when reporting observational studies in conference abstracts. https://www.strobe-statement.org/fileadmin/Strobe/uploads/checklists/STROBE_checklist_conference_abstract_DRAFT.pdf |  |  |  |
| RG127 | Fisher M, Feuerstein G, Howells DW, Hurn PD, Kent TA, Savitz SI, et al. Update of the stroke therapy academic industry roundtable preclinical recommendations. Stroke. 2009;40(6):2244-50. |  |  |  |
| RG128 | Fitchett EJA, Seale AC, Vergnano S, Sharland M, Heath PT, Saha SK, et al. Strengthening the Reporting of Observational Studies in Epidemiology for Newborn Infection (STROBE-NI): an extension of the STROBE statement for neonatal infection research. The Lancet Infectious diseases. 2016;16(10):e202-e13. | STROBE-NI |  |  |
| RG129 | Fletcher R, Ferris L. World Association of Medical Editors (WAME) Editorial Policy and Publication Ethics Committees. Conflict of Interest in Peer-Reviewed Medical Journals2009. Available from: http://www.wame.org/conflict-of-interest-in-peer-reviewed-medical-journals. |  |  |  |
| RG130 | Flores SA, Crepaz N. Quality of study methods in individual- and group-level HIV intervention research: critical reporting elements. AIDS education and prevention : official publication of the International Society for AIDS Education. 2004;16(4):341-52. |  |  |  |
| RG131 | Froud R, Eldridge S, Kovacs F, Breen A, Bolton J, Dunn K, et al. Reporting outcomes of back pain trials: a modified Delphi study. European journal of pain (London, England). 2011;15(10):1068-74. |  |  |  |
| RG132 | Fumagalli D, Bedard PL, Nahleh Z, Michiels S, Sotiriou C, Loi S, et al. A common language in neoadjuvant breast cancer clinical trials: proposals for standard definitions and endpoints. The Lancet Oncology. 2012;13(6):e240-8. |  |  |  |
| RG133 | Gagnier JJ, Boon H, Rochon P, Moher D, Barnes J, Bombardier C. Reporting randomized, controlled trials of herbal interventions: an elaborated CONSORT statement. Annals of internal medicine. 2006;144(5):364-7. | CONSORT Herbal |  |  |
| RG134 | Gagnier JJ, Riley D, Altman DG, Moher D, Sox H, Kienle G. The CARE guidelines: consensus-based clinical case reporting guideline development. Deutsches Arzteblatt international. 2013;110(37):603-8. | CARE |  |  |
| RG135 | Gallo V, Egger M, McCormack V, Farmer PB, Ioannidis JP, Kirsch-Volders M, et al. STrengthening the reporting of OBservational studies in Epidemiology-Molecular Epidemiology (STROBE-ME): an extension of the STROBE statement. European journal of epidemiology. 2011;26(10):797-810. | STROBE-ME |  |  |
| RG136 | Gamble C, Krishan A, Stocken D, Lewis S, Juszczak E, Dore C, et al. Guidelines for the Content of Statistical Analysis Plans in Clinical Trials. Jama. 2017;318(23):2337-43. |  |  |  |
| RG137 | Gardner IA, Nielsen SS, Whittington RJ, Collins MT, Bakker D, Harris B, et al. Consensus-based reporting standards for diagnostic test accuracy studies for paratuberculosis in ruminants. Preventive veterinary medicine. 2011;101(1-2):18-34. | STRADAS-paraTB |  |  |
| RG138 | Gershman MD, Staples JE, Bentsi-Enchill AD, Breugelmans JG, Brito GS, Camacho LA, et al. Viscerotropic disease: case definition and guidelines for collection, analysis, and presentation of immunization safety data. Vaccine. 2012;30(33):5038-58. |  |  |  |
| RG139 | Gewandter JS, Eisenach JC, Gross RA, Jensen MP, Keefe FJ, Lee DA, et al. Checklist for the preparation and review of pain clinical trial publications: a pain-specific supplement to CONSORT. PAIN Reports. 2018. |  |  |  |
| RG140 | Gidudu J, Sack DA, Pina M, Hudson MJ, Kohl KS, Bishop P, et al. Diarrhea: case definition and guidelines for collection, analysis, and presentation of immunization safety data. Vaccine. 2011;29(5):1053-71. |  |  |  |
| RG141 | Gidudu JF, Walco GA, Taddio A, Zempsky WT, Halperin SA, Calugar A, et al. Immunization site pain: case definition and guidelines for collection, analysis, and presentation of immunization safety data. Vaccine. 2012;30(30):4558-77. |  |  |  |
| RG142 | Gnanapavan S, Hegen H, Khalil M, Hemmer B, Franciotta D, Hughes S, et al. Guidelines for uniform reporting of body fluid biomarker studies in neurologic disorders. Neurology. 2014;83(13):1210-6. |  |  |  |
| RG143 | Gold MS, Gidudu J, Erlewyn-Lajeunesse M, Law B. Can the Brighton Collaboration case definitions be used to improve the quality of Adverse Event Following Immunization (AEFI) reporting? Anaphylaxis as a case study. Vaccine. 2010;28(28):4487-98. |  |  |  |
| RG144 | Gotta V, Dao K, Rodieux F, Buclin T, Livio F, Pfister M. Guidance to develop individual dose recommendations for patients on chronic hemodialysis. Expert review of clinical pharmacology. 2017;10(7):737-52. |  |  |  |
| RG145 | Gøtzsche PC, Kassirer JP, Woolley KL, Wager E, Jacobs A, Gertel A, et al. What should be done to tackle ghostwriting in the medical literature? PLoS medicine. 2009;6(2):e23-e. |  |  |  |
| RG146 | Gray RJ, Sacks D, Martin LG, Trerotola SO. Reporting standards for percutaneous interventions in dialysis access. Journal of vascular and interventional radiology : JVIR. 2003;14(9 Pt 2):S433-42. |  |  |  |
| RG147 | Groeneweg R, Rubinstein SM, Oostendorp RA, Ostelo RW, van Tulder MW. Guideline for Reporting Interventions on Spinal Manipulative Therapy: Consensus on Interventions Reporting Criteria List for Spinal Manipulative Therapy (CIRCLe SMT). Journal of manipulative and physiological therapeutics. 2017;40(2):61-70. | CIRCLe SMT |  |  |
| RG148 | Guilhot J, Baccarani M, Clark RE, Cervantes F, Guilhot F, Hochhaus A, et al. Definitions, methodological and statistical issues for phase 3 clinical trials in chronic myeloid leukemia: a proposal by the European LeukemiaNet. Blood. 2012;119(25):5963-71. |  |  |  |
| RG149 | Guise JM, Butler ME, Chang C, Viswanathan M, Pigott T, Tugwell P. AHRQ series on complex intervention systematic reviews-paper 6: PRISMA-CI extension statement and checklist. Journal of clinical epidemiology. 2017;90:43-50. | PRISMA-CI |  |  |
| RG150 | Gurgel RK, Jackler RK, Dobie RA, Popelka GR. A new standardized format for reporting hearing outcome in clinical trials. Otolaryngology--head and neck surgery : official journal of American Academy of Otolaryngology-Head and Neck Surgery. 2012;147(5):803-7. |  |  |  |
| RG151 | Hagen NA, Wu JS, Stiles CR. A proposed taxonomy of terms to guide the clinical trial recruitment process. Journal of pain and symptom management. 2010;40(1):102-10. |  |  |  |
| RG152 | Haidet P, Levine RE, Parmelee DX, Crow S, Kennedy F, Kelly PA, et al. Perspective: Guidelines for reporting team-based learning activities in the medical and health sciences education literature. Academic medicine : journal of the Association of American Medical Colleges. 2012;87(3):292-9. |  |  |  |
| RG153 | Hakoum MB, Jouni N, Abou-Jaoude EA, Hasbani DJ, Abou-Jaoude EA, Lopes LC, et al. Characteristics of funding of clinical trials: cross-sectional survey and proposed guidance. BMJ Open. 2017;7(10):e015997. |  |  |  |
| RG154 | Hamilton S, Bernstein AB, Blakey G, Fagan V, Farrow T, Jordan D, et al. Developing the Clarity and Openness in Reporting: E3-based (CORE) Reference user manual for creation of clinical study reports in the era of clinical trial transparency. Research integrity and peer review. 2016;1:4. | CORE |  |  |
| RG155 | Harrington NG, Noar SM. Reporting standards for studies of tailored interventions. Health education research. 2012;27(2):331-42. |  |  |  |
| RG156 | Heidari S, Babor TF, De Castro P, Tort S, Curno M. Sex and Gender Equity in Research: rationale for the SAGER guidelines and recommended use. Research integrity and peer review. 2016;1:2. | SAGER |  |  |
| RG157 | Helmhout PH, Staal JB, Maher CG, Petersen T, Rainville J, Shaw WS. Exercise therapy and low back pain: insights and proposals to improve the design, conduct, and reporting of clinical trials. Spine. 2008;33(16):1782-8. |  |  |  |
| RG158 | Hemming K, Haines TP, Chilton PJ, Girling AJ, Lilford RJ. The stepped wedge cluster randomised trial: rationale, design, analysis, and reporting. BMJ (Clinical research ed). 2015;350:h391. | SW-CRT |  |  |
| RG159 | Higashida RT, Furlan AJ, Roberts H, Tomsick T, Connors B, Barr J, et al. Trial design and reporting standards for intra-arterial cerebral thrombolysis for acute ischemic stroke. Stroke. 2003;34(8):e109-37. |  |  |  |
| RG160 | Higashida RT, Meyers PM, Phatouros CC, Connors JJ, 3rd, Barr JD, Sacks D. Reporting standards for carotid artery angioplasty and stent placement. Stroke. 2004;35(5):e112-34. |  |  |  |
| RG161 | Higgins JPT, Green S, editors. Cochrane Handbook for Systematic Reviews of Interventions Version 5.1.0: The Cochrane Collaboration; 2011. |  |  |  |
| RG162 | Higginson IJ, Evans CJ, Grande G, Preston N, Morgan M, McCrone P, et al. Evaluating complex interventions in End of Life Care: the MORECare Statement on good practice generated by a synthesis of transparent expert consultations and systematic reviews. 2013;11(1):111. | MORECare |  |  |
| RG163 | Hlatky MA, Greenland P, Arnett DK, Ballantyne CM, Criqui MH, Elkind MS, et al. Criteria for evaluation of novel markers of cardiovascular risk: a scientific statement from the American Heart Association. Circulation. 2009;119(17):2408-16. |  |  |  |
| RG164 | Hoffmann TC, Glasziou PP, Boutron I, Milne R, Perera R, Moher D, et al. Better reporting of interventions: template for intervention description and replication (TIDieR) checklist and guide. BMJ (Clinical research ed). 2014;348:g1687. | TIDieR |  |  |
| RG165 | Hollander JE, Blomkalns AL, Brogan GX, Diercks DB, Field JM, Garvey JL, et al. Standardized reporting guidelines for studies evaluating risk stratification of ED patients with potential acute coronary syndromes. Academic emergency medicine : official journal of the Society for Academic Emergency Medicine. 2004;11(12):1331-40. |  |  |  |
| RG166 | Hollenbach JA, Mack SJ, Gourraud PA, Single RM, Maiers M, Middleton D, et al. A community standard for immunogenomic data reporting and analysis: proposal for a STrengthening the REporting of Immunogenomic Studies statement. Tissue antigens. 2011;78(5):333-44. | STREIS |  |  |
| RG167 | Holtfreter B, Albandar JM, Dietrich T, Dye BA, Eaton KA, Eke PI, et al. Standards for reporting chronic periodontitis prevalence and severity in epidemiologic studies: Proposed standards from the Joint EU/USA Periodontal Epidemiology Working Group. Journal of clinical periodontology. 2015;42(5):407-12. |  |  |  |
| RG168 | Hooijmans CR, Leenaars M, Ritskes-Hoitinga M. A gold standard publication checklist to improve the quality of animal studies, to fully integrate the Three Rs, and to make systematic reviews more feasible. Alternatives to laboratory animals : ATLA. 2010;38(2):167-82. |  |  |  |
| RG169 | Hopewell S, Clarke M, Moher D, Wager E, Middleton P, Altman DG, et al. CONSORT for reporting randomised trials in journal and conference abstracts. Lancet (London, England). 2008;371(9609):281-3. | CONSORT for abstracts |  |  |
| RG170 | Horby PW, Laurie KL, Cowling BJ, Engelhardt OG, Sturm-Ramirez K, Sanchez JL, et al. CONSISE statement on the reporting of Seroepidemiologic Studies for influenza (ROSES-I statement): an extension of the STROBE statement. Influenza and other respiratory viruses. 2017;11(1):2-14. | ROSES-I |  |  |
| RG171 | Horstkotte D, Lengyel M, Mistiaen WP, Piper C, Voller H. Recommendations for reporting morbid events after heart valve surgery. The Journal of heart valve disease. 2005;14(1):1-7. |  |  |  |
| RG172 | Howley L, Szauter K, Perkowski L, Clifton M, McNaughton N. Quality of standardised patient research reports in the medical education literature: review and recommendations. Medical education. 2008;42(4):350-8. |  |  |  |
| RG173 | Hrynaszkiewicz I, Norton ML, Vickers AJ, Altman DG. Preparing raw clinical data for publication: guidance for journal editors, authors, and peer reviewers. BMJ (Clinical research ed). 2010;340:c181. |  |  |  |
| RG174 | Humar A, Michaels M. American Society of Transplantation recommendations for screening, monitoring and reporting of infectious complications in immunosuppression trials in recipients of organ transplantation. American journal of transplantation : official journal of the American Society of Transplantation and the American Society of Transplant Surgeons. 2006;6(2):262-74. |  |  |  |
| RG175 | Hundley WG, Bluemke D, Bogaert JG, Friedrich MG, Higgins CB, Lawson MA, et al. Society for Cardiovascular Magnetic Resonance guidelines for reporting cardiovascular magnetic resonance examinations. Journal of cardiovascular magnetic resonance : official journal of the Society for Cardiovascular Magnetic Resonance. 2009;11:5. |  |  |  |
| RG176 | Husereau D, Drummond M, Petrou S, Carswell C, Moher D, Greenberg D, et al. Consolidated Health Economic Evaluation Reporting Standards (CHEERS) statement. Value in health : the journal of the International Society for Pharmacoeconomics and Outcomes Research. 2013;16(2):e1-5. | CHEERS |  |  |
| RG177 | Hutton B, Salanti G, Caldwell DM, Chaimani A, Schmid CH, Cameron C, et al. The PRISMA extension statement for reporting of systematic reviews incorporating network meta-analyses of health care interventions: checklist and explanations. Annals of internal medicine. 2015;162(11):777-84. |  |  |  |
| RG178 | ICMJE: Uniform Format for Disclosure of Competing Interests in ICMJE Journals. 2010 [updated July 2010. Available from: http://www.icmje.org/coi_disclosure.pdf. Accessed 12 June 2019. |  |  |  |
| RG179 | Idris AH, Becker LB, Ornato JP, Hedges JR, Bircher NG, Chandra NC, et al. Utstein-style guidelines for uniform reporting of laboratory CPR research. A statement for healthcare professionals from a task force of the American Heart Association, the American College of Emergency Physicians, the American College of Cardiology, the European Resuscitation Council, the Heart and Stroke Foundation of Canada, the Institute of Critical Care Medicine, the Safar Center for Resuscitation Research, and the Society for Academic Emergency Medicine. Writing Group. Circulation. 1996;94(9):2324-36. |  |  |  |
| RG180 | Iglesias CP, Thompson A, Rogowski WH, Payne K. Reporting Guidelines for the Use of Expert Judgement in Model-Based Economic Evaluations. PharmacoEconomics. 2016;34(11):1161-72. |  |  |  |
| RG181 | Institute of Medicine Committee on Standards for Systematic Reviews of Comparative Effectiveness R. Finding what works in health care: standards for systematic reviews. In: Eden J, Levit L, Berg A, Morton S, editors. Finding What Works in Health Care: Standards for Systematic Reviews. Washington (DC): National Academies Press (US) Copyright 2011 by the National Academy of Sciences. All rights reserved.; 2011. |  |  |  |
| RG182 | International Society for Medical Publication Professionals Code of Ethics. Available from: http://www.ismpp.org/ethics. |  |  |  |
| RG183 | Ioannidis JP, Evans SJ, Gotzsche PC, O'Neill RT, Altman DG, Schulz K, et al. Better reporting of harms in randomized trials: an extension of the CONSORT statement. Annals of internal medicine. 2004;141(10):781-8. | CONSORT Harms |  |  |
| RG184 | Jabs DA. Improving the reporting of clinical case series. American journal of ophthalmology. 2005;139(5):900-5. |  |  |  |
| RG185 | Jabs DA, Nussenblatt RB, Rosenbaum JT. Standardization of uveitis nomenclature for reporting clinical data. Results of the First International Workshop. American journal of ophthalmology. 2005;140(3):509-16. |  |  |  |
| RG186 | Jackson A, Marks LB, Bentzen SM, Eisbruch A, Yorke ED, Ten Haken RK, et al. The lessons of QUANTEC: recommendations for reporting and gathering data on dose-volume dependencies of treatment outcome. International journal of radiation oncology, biology, physics. 2010;76(3 Suppl):S155-60. |  |  |  |
| RG187 | Jackson DL. Reporting results of latent growth modeling and multilevel modeling analyses: some recommendations for rehabilitation psychology. Rehabilitation psychology. 2010;55(3):272-85. |  |  |  |
| RG188 | Janssens AC, Ioannidis JP, van Duijn CM, Little J, Khoury MJ. Strengthening the reporting of genetic risk prediction studies: The GRIPS Statement. Annals of internal medicine. 2011;154(6):421-5. |  |  |  |
| RG189 | Jason LA, Unger ER, Dimitrakoff JD, Fagin AP, Houghton M, Cook DB, et al. Minimum data elements for research reports on CFS. Brain, behavior, and immunity. 2012;26(3):401-6. |  |  |  |
| RG190 | Jayaraman MV, Meyers PM, Derdeyn CP, Fraser JF, Hirsch JA, Hussain MS, et al. Reporting standards for angiographic evaluation and endovascular treatment of cerebral arteriovenous malformations. Journal of neurointerventional surgery. 2012;4(5):325-30. |  |  |  |
| RG191 | Jenkins PA, Carroll JD. How to report low-level laser therapy (LLLT)/photomedicine dose and beam parameters in clinical and laboratory studies. Photomedicine and laser surgery. 2011;29(12):785-7. |  |  |  |
| RG192 | Junger S, Payne SA, Brine J, Radbruch L, Brearley SG. Guidance on Conducting and REporting DElphi Studies (CREDES) in palliative care: Recommendations based on a methodological systematic review. Palliative medicine. 2017;31(8):684-706. | CREDES |  |  |
| RG193 | Kable AK, Pich J, Maslin-Prothero SE. A structured approach to documenting a search strategy for publication: a 12 step guideline for authors. Nurse education today. 2012;32(8):878-86. |  |  |  |
| RG194 | Kahn MG, Brown JS, Chun AT, Davidson BN, Meeker D, Ryan PB, et al. Transparent reporting of data quality in distributed data networks. EGEMS (Washington, DC). 2015;3(1):1052. |  |  |  |
| RG195 | Kalil AC, Mattei J, Florescu DF, Sun J, Kalil RS. Recommendations for the assessment and reporting of multivariable logistic regression in transplantation literature. American journal of transplantation : official journal of the American Society of Transplantation and the American Society of Transplant Surgeons. 2010;10(7):1686-94. |  |  |  |
| RG196 | Kaloupek DG, Chard KM, Freed MC, Peterson AL, Riggs DS, Stein MB, et al. Common data elements for posttraumatic stress disorder research. Archives of physical medicine and rehabilitation. 2010;91(11):1684-91. |  |  |  |
| RG197 | Kanji S, Hayes M, Ling A, Shamseer L, Chant C, Edwards DJ, et al. Reporting Guidelines for Clinical Pharmacokinetic Studies: The ClinPK Statement. Clinical pharmacokinetics. 2015;54(7):783-95. | ClinPK |  |  |
| RG198 | Karmy-Jones R, Ferrigno L, Teso D, Long WB, 3rd, Shackford S. Endovascular repair compared with operative repair of traumatic rupture of the thoracic aorta: a nonsystematic review and a plea for trauma-specific reporting guidelines. The Journal of trauma. 2011;71(4):1059-72. |  |  |  |
| RG199 | Kearney HM, Thorland EC, Brown KK, Quintero-Rivera F, South ST. American College of Medical Genetics standards and guidelines for interpretation and reporting of postnatal constitutional copy number variants. Genetics in medicine : official journal of the American College of Medical Genetics. 2011;13(7):680-5. |  |  |  |
| RG200 | Kearon C, Iorio A, Palareti G. Risk of recurrent venous thromboembolism after stopping treatment in cohort studies: recommendation for acceptable rates and standardized reporting. Journal of thrombosis and haemostasis : JTH. 2010;8(10):2313-5. |  |  |  |
| RG201 | Keith MG, Tay L, Harms PD. Systems Perspective of Amazon Mechanical Turk for Organizational Research: Review and Recommendations. Frontiers in psychology. 2017;8:1359. |  |  |  |
| RG202 | Kelley K, Clark B, Brown V, Sitzia J. Good practice in the conduct and reporting of survey research. International journal for quality in health care : journal of the International Society for Quality in Health Care. 2003;15(3):261-6. |  |  |  |
| RG203 | Kelly WN, Arellano FM, Barnes J, Bergman U, Edwards IR, Fernandez AM, et al. Guidelines for submitting adverse event reports for publication. Pharmacoepidemiology and drug safety. 2007;16(5):581-7. |  |  |  |
| RG204 | Kempen JH. Appropriate use and reporting of uncontrolled case series in the medical literature. American journal of ophthalmology. 2011;151(1):7-10.e1. |  |  |  |
| RG205 | Kent DM, Rothwell PM, Ioannidis JP, Altman DG, Hayward RA. Assessing and reporting heterogeneity in treatment effects in clinical trials: a proposal. Trials. 2010;11:85. |  |  |  |
| RG206 | Kerr KF, Meisner A, Thiessen-Philbrook H, Coca SG, Parikh CR. RiGoR: reporting guidelines to address common sources of bias in risk model development. Biomarker research. 2015;3(1):2-. | RiGoR |  |  |
| RG207 | Keyerleber MA, McEntee MC, Farrelly J, Podgorsak M. Completeness of reporting of radiation therapy planning, dose, and delivery in veterinary radiation oncology manuscripts from 2005 to 2010. Veterinary radiology & ultrasound : the official journal of the American College of Veterinary Radiology and the International Veterinary Radiology Association. 2012;53(2):221-30. |  |  |  |
| RG208 | Khanal S, Burgon J, Leonard S, Griffiths M, Eddowes LA. Recommendations for the Improved Effectiveness and Reporting of Telemedicine Programs in Developing Countries: Results of a Systematic Literature Review. Telemedicine journal and e-health : the official journal of the American Telemedicine Association. 2015;21(11):903-15. |  |  |  |
| RG209 | Kilkenny C, Browne WJ, Cuthill IC, Emerson M, Altman DG. Improving bioscience research reporting: the ARRIVE guidelines for reporting animal research. PLoS biology. 2010;8(6):e1000412. | ARRIVE |  |  |
| RG210 | Kirkham JJ, Gorst S, Altman DG, Blazeby JM, Clarke M, Devane D, et al. Core Outcome Set-STAndards for Reporting: The COS-STAR Statement. PLoS medicine. 2016;13(10):e1002148. | COS-STAR |  |  |
| RG211 | Kostoulas P, Nielsen SS, Branscum AJ, Johnson WO, Dendukuri N, Dhand NK, et al. STARD-BLCM: Standards for the Reporting of Diagnostic accuracy studies that use Bayesian Latent Class Models. Preventive veterinary medicine. 2017;138:37-47. | STARD-BLCM |  |  |
| RG212 | Kottner J, Audige L, Brorson S, Donner A, Gajewski BJ, Hrobjartsson A, et al. Guidelines for Reporting Reliability and Agreement Studies (GRRAS) were proposed. Journal of clinical epidemiology. 2011;64(1):96-106. | GRRAS |  |  |
| RG213 | Lachat C, Hawwash D, Ocke MC, Berg C, Forsum E, Hornell A, et al. Strengthening the Reporting of Observational Studies in Epidemiology-Nutritional Epidemiology (STROBE-nut): An Extension of the STROBE Statement. PLoS medicine. 2016;13(6):e1002036. | STROBE-nut |  |  |
| RG214 | Landis RC, Arrowsmith JE, Baker RA, de Somer F, Dobkowski WB, Fisher G, et al. Consensus statement: Defining minimal criteria for reporting the systemic inflammatory response to cardiopulmonary bypass. The heart surgery forum. 2008;11(5):E316-22. |  |  |  |
| RG215 | Landis SC, Amara SG, Asadullah K, Austin CP, Blumenstein R, Bradley EW, et al. A call for transparent reporting to optimize the predictive value of preclinical research. Nature. 2012;490(7419):187-91. |  |  |  |
| RG216 | Lang TA, Altman DG. Basic statistical reporting for articles published in biomedical journals: the "Statistical Analyses and Methods in the Published Literature" or the SAMPL Guidelines. International journal of nursing studies. 2015;52(1):5-9. | SAMPL |  |  |
| RG217 | Lang TA, Talerico C, Siontis GCM. Documenting clinical and laboratory images in publications: the CLIP principles. Chest. 2012;141(6):1626-32. | CLIP principles |  |  |
| RG218 | Langan SM, Schmidt SA, Wing K, Ehrenstein V, Nicholls SG, Filion KB, et al. The reporting of studies conducted using observational routinely collected health data statement for pharmacoepidemiology (RECORD-PE). 2018;363. | RECORD-PE |  |  |
| RG219 | Langhelle A, Nolan J, Herlitz J, Castren M, Wenzel V, Soreide E, et al. Recommended guidelines for reviewing, reporting, and conducting research on post-resuscitation care: the Utstein style. Resuscitation. 2005;66(3):271-83. |  |  |  |
| RG220 | Lassmann M, Chiesa C, Flux G, Bardies M. EANM Dosimetry Committee guidance document: good practice of clinical dosimetry reporting. European journal of nuclear medicine and molecular imaging. 2011;38(1):192-200. |  |  |  |
| RG221 | Launay E, Cohen JF, Bossuyt PM, Buekens P, Deeks J, Dye T, et al. Reporting studies on time to diagnosis: proposal of a guideline by an international panel (REST). BMC medicine. 2016;14(1):146. | REST |  |  |
| RG222 | Lavergne V, Ouellet G, Bouchard J, Galvao T, Kielstein JT, Roberts DM, et al. Guidelines for reporting case studies on extracorporeal treatments in poisonings: methodology. Seminars in dialysis. 2014;27(4):407-14. |  |  |  |
| RG223 | Lee CW, Chi KN. The standard of reporting of health-related quality of life in clinical cancer trials. Journal of Clinical Epidemiology. 2000;53(5):451-8. |  |  |  |
| RG224 | Leech NL, Onwuegbuzie AJ. Guidelines for Conducting and Reporting Mixed Research in the Field of Counseling and Beyond. Journal of Counseling & Development. 2010;88(1):61-9. |  |  |  |
| RG225 | Legro RS, Wu X, Barnhart KT, Farquhar C, Fauser BC, Mol B. Improving the reporting of clinical trials of infertility treatments (IMPRINT): modifying the CONSORT statementdaggerdouble dagger. Human reproduction (Oxford, England). 2014;29(10):2075-82. | IMPRINT |  |  |
| RG226 | Leipsic J, Abbara S, Achenbach S, Cury R, Earls JP, Mancini GJ, et al. SCCT guidelines for the interpretation and reporting of coronary CT angiography: a report of the Society of Cardiovascular Computed Tomography Guidelines Committee. Journal of cardiovascular computed tomography. 2014;8(5):342-58. |  |  |  |
| RG227 | Lineberry N, Berlin JA, Mansi B, Glasser S, Berkwits M, Klem C, et al. Recommendations to improve adverse event reporting in clinical trial publications: a joint pharmaceutical industry/journal editor perspective. 2016;355:i5078. |  |  |  |
| RG228 | Little J, Higgins JP, Ioannidis JP, Moher D, Gagnon F, von Elm E, et al. STrengthening the REporting of Genetic Association Studies (STREGA)--an extension of the STROBE statement. Genetic epidemiology. 2009;33(7):581-98. | STREGA |  |  |
| RG229 | Little J, Higgins JPT, editors. The HuGENet™ HuGE Review Handbook, version 1.0: Centers for Disease Control and Prevention; 2006. |  |  |  |
| RG230 | Luo W, Phung D, Tran T, Gupta S, Rana S, Karmakar C, et al. Guidelines for Developing and Reporting Machine Learning Predictive Models in Biomedical Research: A Multidisciplinary View. Journal of medical Internet research. 2016;18(12):e323. |  |  |  |
| RG231 | Macleod MR, Fisher M, O'Collins V, Sena ES, Dirnagl U, Bath PM, et al. Good laboratory practice: preventing introduction of bias at the bench. Stroke. 2009;40(3):e50-2. |  |  |  |
| RG232 | MacPherson H, Altman DG, Hammerschlag R, Youping L, Taixiang W, White A, et al. Revised STandards for Reporting Interventions in Clinical Trials of Acupuncture (STRICTA): extending the CONSORT statement. PLoS medicine. 2010;7(6):e1000261. | STRICTA |  |  |
| RG233 | Madurasinghe VW. Guidelines for reporting embedded recruitment trials. Trials. 2016;17:27. |  |  |  |
| RG234 | Malterud K. Qualitative research: standards, challenges, and guidelines. Lancet (London, England). 2001;358(9280):483-8. |  |  |  |
| RG235 | Matcham J, Julious S, Pyke S, O'Kelly M, Todd S, Seldrup J, et al. Proposed best practice for statisticians in the reporting and publication of pharmaceutical industry-sponsored clinical trials. Pharmaceutical statistics. 2011;10(1):70-3. |  |  |  |
| RG236 | McInnes MDF, Moher D, Thombs BD, McGrath TA, Bossuyt PM, Clifford T, et al. Preferred Reporting Items for a Systematic Review and Meta-analysis of Diagnostic Test Accuracy Studies: The PRISMA-DTA Statement. Jama. 2018;319(4):388-96. | PRISMA-DTA |  |  |
| RG237 | McShane LM, Altman DG, Sauerbrei W, Taube SE, Gion M, Clark GM. REporting recommendations for tumour MARKer prognostic studies (REMARK). British journal of cancer. 2005;93(4):387-91. | REMARK |  |  |
| RG238 | Melby MK, Sievert LL, Anderson D, Obermeyer CM. Overview of methods used in cross-cultural comparisons of menopausal symptoms and their determinants: Guidelines for Strengthening the Reporting of Menopause and Aging (STROMA) studies. Maturitas. 2011;70(2):99-109. | STROMA |  |  |
| RG239 | Messiou C, Bonvalot S, Gronchi A, Vanel D, Meyer M, Robinson P, et al. Evaluation of response after pre-operative radiotherapy in soft tissue sarcomas; the European Organisation for Research and Treatment of Cancer-Soft Tissue and Bone Sarcoma Group (EORTC-STBSG) and Imaging Group recommendations for radiological examination and reporting with an emphasis on magnetic resonance imaging. European journal of cancer (Oxford, England : 1990). 2016;56:37-44. |  |  |  |
| RG240 | Meyer E, Delaney M, Lin Y, Morris A, Pavenski K, Tinmouth A, et al. A reporting guideline for clinical platelet transfusion studies from the BEST Collaborative. Transfusion. 2013;53(6):1328-34. |  |  |  |
| RG241 | Meyers PM, Schumacher HC, Higashida RT, Derdeyn CP, Nesbit GM, Sacks D, et al. Reporting standards for endovascular repair of saccular intracranial cerebral aneurysms. Journal of vascular and interventional radiology : JVIR. 2009;20(7 Suppl):S435-50. |  |  |  |
| RG242 | Mintz GS, Nissen SE, Anderson WD, Bailey SR, Erbel R, Fitzgerald PJ, et al. American College of Cardiology Clinical Expert Consensus Document on Standards for Acquisition, Measurement and Reporting of Intravascular Ultrasound Studies (IVUS). A report of the American College of Cardiology Task Force on Clinical Expert Consensus Documents. Journal of the American College of Cardiology. 2001;37(5):1478-92. |  |  |  |
| RG243 | Mischak H, Allmaier G, Apweiler R, Attwood T, Baumann M, Benigni A, et al. Recommendations for biomarker identification and qualification in clinical proteomics. Science translational medicine. 2010;2(46):46ps2. |  |  |  |
| RG244 | Mistry P, Dunn JA, Marshall A. A literature review of applied adaptive design methodology within the field of oncology in randomised controlled trials and a proposed extension to the CONSORT guidelines. BMC Medical Research Methodology. 2017;17(1):108. |  |  |  |
| RG245 | Moberg-Mogren E, Nelson DL. Evaluating the quality of reporting occupational therapy randomized controlled trials by expanding the CONSORT criteria. The American journal of occupational therapy: official publication of the American Occupational Therapy Association. 2006;60(2):226-35. |  |  |  |
| RG246 | Moher D, Liberati A, Tetzlaff J, Altman DG. Preferred reporting items for systematic reviews and meta-analyses: the PRISMA statement. Annals of internal medicine. 2009;151(4):264-9, w64. | PRISMA |  |  |
| RG247 | Moher D, Shamseer L, Clarke M, Ghersi D, Liberati A, Petticrew M, et al. Preferred reporting items for systematic review and meta-analysis protocols (PRISMA-P) 2015 statement. Systematic reviews. 2015;4:1. | PRISMA-P |  |  |
| RG248 | Möhler R, Köpke S, Meyer GJT. Criteria for Reporting the Development and Evaluation of Complex Interventions in healthcare: revised guideline (CReDECI 2). 2015;16(1):204. | CReDECI 2 |  |  |
| RG249 | Monks T, Currie CSM, Onggo BS, Robinson S, Kunc M, Taylor SJE. Strengthening the reporting of empirical simulation studies: Introducing the STRESS guidelines. Journal of Simulation. 2018:1-13. | STRESS |  |  |
| RG250 | Montgomery P, Grant S, Mayo-Wilson E, Macdonald G, Michie S, Hopewell S, et al. Reporting randomised trials of social and psychological interventions: the CONSORT-SPI 2018 Extension. 2018;19(1):407. | CONSORT-SPI 2018 |  |  |
| RG251 | Moons KG, Altman DG, Reitsma JB, Ioannidis JP, Macaskill P, Steyerberg EW, et al. Transparent Reporting of a multivariable prediction model for Individual Prognosis or Diagnosis (TRIPOD): explanation and elaboration. Annals of internal medicine. 2015;162(1):W1-73. | TRIPOD |  |  |
| RG252 | Moore CM, Giganti F, Albertsen P, Allen C, Bangma C, Briganti A, et al. Reporting Magnetic Resonance Imaging in Men on Active Surveillance for Prostate Cancer: The PRECISE Recommendations-A Report of a European School of Oncology Task Force. European urology. 2017;71(4):648-55. | PRECISE |  |  |
| RG253 | Moore CM, Kasivisvanathan V, Eggener S, Emberton M, Futterer JJ, Gill IS, et al. Standards of reporting for MRI-targeted biopsy studies (START) of the prostate: recommendations from an International Working Group. European urology. 2013;64(4):544-52. | START |  |  |
| RG254 | Moore HM, Kelly AB, Jewell SD, McShane LM, Clark DP, Greenspan R, et al. Biospecimen reporting for improved study quality (BRISQ). Cancer cytopathology. 2011;119(2):92-101. | BRISQ |  |  |
| RG255 | Moss F, Thomson R. A new structure for quality improvement reports. Quality & safety in health care. 2004;13(1):6-7. |  |  |  |
| RG256 | Muller VI, Cieslik EC, Laird AR, Fox PT, Radua J, Mataix-Cols D, et al. Ten simple rules for neuroimaging meta-analysis. Neuroscience and biobehavioral reviews. 2018;84:151-61. |  |  |  |
| RG257 | Munk N, Boulanger K. Adaptation of the CARE Guidelines for Therapeutic Massage and Bodywork Publications: Efforts To Improve the Impact of Case Reports. International journal of therapeutic massage & bodywork. 2014;7(3):32-40. |  |  |  |
| RG258 | Murad MH, Wang Z. Guidelines for reporting meta-epidemiological methodology research. Evidence-based medicine. 2017;22(4):139-42. |  |  |  |
| RG259 | Murray IR, Geeslin AG, Goudie EB, Petrigliano FA, LaPrade RF. Minimum Information for Studies Evaluating Biologics in Orthopaedics (MIBO): Platelet-Rich Plasma and Mesenchymal Stem Cells. The Journal of bone and joint surgery American volume. 2017;99(10):809-19. | MIBO |  |  |
| RG260 | Nedeltchev K, Pattynama PM, Biaminoo G, Diehm N, Jaff MR, Hopkins LN, et al. Standardized definitions and clinical endpoints in carotid artery and supra-aortic trunk revascularization trials. Catheterization and cardiovascular interventions : official journal of the Society for Cardiac Angiography & Interventions. 2010;76(3):333-44. |  |  |  |
| RG261 | Newton PN, Lee SJ, Goodman C, Fernandez FM, Yeung S, Phanouvong S, et al. Guidelines for field surveys of the quality of medicines: a proposal. PLoS medicine. 2009;6(3):e52. |  |  |  |
| RG262 | Nichols TE, Das S, Eickhoff SB, Evans AC, Glatard T, Hanke M, et al. Best practices in data analysis and sharing in neuroimaging using MRI. Nature neuroscience. 2017;20(3):299-303. |  |  |  |
| RG263 | Nicholson A, Berger K, Bohn R, Carcao M, Fischer K, Gringeri A, et al. Recommendations for reporting economic evaluations of haemophilia prophylaxis: a nominal groups consensus statement on behalf of the Economics Expert Working Group of The International Prophylaxis Study Group. Haemophilia : the official journal of the World Federation of Hemophilia. 2008;14(1):127-32. |  |  |  |
| RG264 | Niederstadt C, Droste S. Reporting and presenting information retrieval processes: the need for optimizing common practice in health technology assessment. International journal of technology assessment in health care. 2010;26(4):450-7. |  |  |  |
| RG265 | Noel-Storr AH, McCleery JM, Richard E, Ritchie CW, Flicker L, Cullum SJ, et al. Reporting standards for studies of diagnostic test accuracy in dementia: The STARDdem Initiative. Neurology. 2014;83(4):364-73. | STARDdem |  |  |
| RG266 | Nothacker M, Stokes T, Shaw B, Lindsay P, Sipila R, Follmann M, et al. Reporting standards for guideline-based performance measures. Implementation science : IS. 2016;11:6. |  |  |  |
| RG267 | Nuijten MJ, Pronk MH, Brorens MJ, Hekster YA, Lockefeer JH, de Smet PA, et al. Reporting format for economic evaluation. Part II: Focus on modelling studies. PharmacoEconomics. 1998;14(3):259-68. |  |  |  |
| RG268 | O'Brien BC, Harris IB, Beckman TJ, Reed DA, Cook DA. Standards for reporting qualitative research: a synthesis of recommendations. Academic medicine : journal of the Association of American Medical Colleges. 2014;89(9):1245-51. | SRQR |  |  |
| RG269 | O'Cathain A, Murphy E, Nicholl J. The quality of mixed methods studies in health services research. Journal of health services research & policy. 2008;13(2):92-8. |  |  |  |
| RG270 | O'Connor AM, Sargeant JM, Gardner IA, Dickson JS, Torrence ME, Dewey CE, et al. The REFLECT statement: methods and processes of creating reporting guidelines for randomized controlled trials for livestock and food safety by modifying the CONSORT statement. Zoonoses and public health. 2010;57(2):95-104. | REFLECT |  |  |
| RG271 | Ogrinc G, Davies L, Goodman D, Batalden P, Davidoff F, Stevens D. SQUIRE 2.0 (Standards for QUality Improvement Reporting Excellence): revised publication guidelines from a detailed consensus process. BMJ quality & safety. 2016;25(12):986-92. | SQUIRE 2.0 |  |  |
| RG272 | Olson SH, Voigt LF, Begg CB, Weiss NS. Reporting participation in case-control studies. Epidemiology (Cambridge, Mass). 2002;13(2):123-6. |  |  |  |
| RG273 | Omulo S, Thumbi SM, Njenga MK, Call DR. A review of 40 years of enteric antimicrobial resistance research in Eastern Africa: what can be done better? Antimicrobial resistance and infection control. 2015;4:1. |  |  |  |
| RG274 | Padhani AR, Lecouvet FE, Tunariu N, Koh DM, De Keyzer F, Collins DJ, et al. METastasis Reporting and Data System for Prostate Cancer: Practical Guidelines for Acquisition, Interpretation, and Reporting of Whole-body Magnetic Resonance Imaging-based Evaluations of Multiorgan Involvement in Advanced Prostate Cancer. European urology. 2017;71(1):81-92. | MET-RADS-P |  |  |
| RG275 | Page P, Hoogenboom B, Voight M. Improving the reporting of therapeutic exercise interventions in rehabilitation research. International journal of sports physical therapy. 2017;12(2):297-304. |  |  |  |
| RG276 | Pandis N, Chung B, Scherer RW, Elbourne D, Altman DG. CONSORT 2010 statement: extension checklist for reporting within person randomised trials. BMJ (Clinical research ed). 2017;357:j2835. |  |  |  |
| RG277 | Pandis N, Fleming PS, Hopewell S, Altman DG. The CONSORT Statement: Application within and adaptations for orthodontic trials. American journal of orthodontics and dentofacial orthopedics : official publication of the American Association of Orthodontists, its constituent societies, and the American Board of Orthodontics. 2015;147(6):663-79. |  |  |  |
| RG278 | Parson W, Roewer L. Publication of population data of linearly inherited DNA markers in the International Journal of Legal Medicine. International journal of legal medicine. 2010;124(5):505-9. |  |  |  |
| RG279 | Patricio M, Juliao M, Fareleira F, Young M, Norman G, Vaz Carneiro A. A comprehensive checklist for reporting the use of OSCEs. Medical teacher. 2009;31(2):112-24. |  |  |  |
| RG280 | Peters JL, Sutton AJ, Jones DR, Rushton L, Abrams KR. A systematic review of systematic reviews and meta-analyses of animal experiments with guidelines for reporting. Journal of environmental science and health Part B, Pesticides, food contaminants, and agricultural wastes. 2006;41(7):1245-58. |  |  |  |
| RG281 | Petrou S, Gray A. Economic evaluation using decision analytical modelling: design, conduct, analysis, and reporting. BMJ (Clinical research ed). 2011;342:d1766. |  |  |  |
| RG282 | Petrou S, Gray A. Economic evaluation alongside randomised controlled trials: design, conduct, analysis, and reporting. BMJ (Clinical research ed). 2011;342:d1548. |  |  |  |
| RG283 | Petrou S, Rivero-Arias O, Dakin H, Longworth L, Oppe M, Froud R, et al. Preferred Reporting Items for Studies Mapping onto Preference-Based Outcome Measures: The MAPS Statement. PharmacoEconomics. 2015;33(10):985-91. | MAPS |  |  |
| RG284 | Phillips AC, Lewis LK, McEvoy MP, Galipeau J, Glasziou P, Moher D, et al. Development and validation of the guideline for reporting evidence-based practice educational interventions and teaching (GREET). 2016;16(1):237. | GREET |  |  |
| RG285 | Piaggio G, Elbourne DR, Pocock SJ, Evans SJ, Altman DG. Reporting of noninferiority and equivalence randomized trials: extension of the CONSORT 2010 statement. Jama. 2012;308(24):2594-604. | CONSORT Non-inferiority |  |  |
| RG286 | Pinnock H, Barwick M, Carpenter CR, Eldridge S, Grandes G, Griffiths CJ, et al. Standards for Reporting Implementation Studies (StaRI) Statement. BMJ (Clinical research ed). 2017;356:i6795. | STARI |  |  |
| RG287 | Pocock SJ, Travison TG, Wruck LM. Figures in clinical trial reports: current practice & scope for improvement. Trials. 2007;8:36. |  |  |  |
| RG288 | Poldrack RA, Fletcher PC, Henson RN, Worsley KJ, Brett M, Nichols TE. Guidelines for reporting an fMRI study. NeuroImage. 2008;40(2):409-14. |  |  |  |
| RG289 | Portell M, Anguera MT, Chacón-Moscoso S, Sanduvete-Chaves S. Guidelines for reporting evaluations based on observational methodology. Psicothema. 2015;27(3):283-9. | GREOM |  |  |
| RG290 | Provenzano E, Bossuyt V, Viale G, Cameron D, Badve S, Denkert C, et al. Standardization of pathologic evaluation and reporting of postneoadjuvant specimens in clinical trials of breast cancer: recommendations from an international working group. Modern pathology : an official journal of the United States and Canadian Academy of Pathology, Inc. 2015;28(9):1185-201. |  |  |  |
| RG291 | Puhan MA, ter Riet G, Eichler K, Steurer J, Bachmann LM. More medical journals should inform their contributors about three key principles of graph construction. Journal of Clinical Epidemiology. 2006;59(10):1017.e1-.e8. |  |  |  |
| RG292 | Quintana DS, Alvares GA, Heathers JA. Guidelines for Reporting Articles on Psychiatry and Heart rate variability (GRAPH): recommendations to advance research communication. Translational psychiatry. 2016;6:e803. | GRAPH |  |  |
| RG293 | Radvany MG, Murphy KJ, Millward SF, Barr JD, Clark TW, Halin NJ, et al. Research reporting standards for percutaneous vertebral augmentation. Journal of vascular and interventional radiology : JVIR. 2009;20(10):1279-86. |  |  |  |
| RG294 | Rajkumar SV, Harousseau JL, Durie B, Anderson KC, Dimopoulos M, Kyle R, et al. Consensus recommendations for the uniform reporting of clinical trials: report of the International Myeloma Workshop Consensus Panel 1. Blood. 2011;117(18):4691-5. |  |  |  |
| RG295 | Ramsey S, Willke R, Briggs A, Brown R, Buxton M, Chawla A, et al. Good research practices for cost-effectiveness analysis alongside clinical trials: the ISPOR RCT-CEA Task Force report. Value in health : the journal of the International Society for Pharmacoeconomics and Outcomes Research. 2005;8(5):521-33. |  |  |  |
| RG296 | Rao SV, Eikelboom J, Steg PG, Lincoff AM, Weintraub WS, Bassand JP, et al. Standardized reporting of bleeding complications for clinical investigations in acute coronary syndromes: a proposal from the academic bleeding consensus (ABC) multidisciplinary working group. American heart journal. 2009;158(6):881-6.e1. |  |  |  |
| RG297 | Rauch F, Sievanen H, Boonen S, Cardinale M, Degens H, Felsenberg D, et al. Reporting whole-body vibration intervention studies: recommendations of the International Society of Musculoskeletal and Neuronal Interactions. Journal of musculoskeletal & neuronal interactions. 2010;10(3):193-8. |  |  |  |
| RG298 | Reeve BB, McFatrich M, Pinheiro LC, Weaver MS, Sung L, Withycombe JS, et al. Eliciting the child's voice in adverse event reporting in oncology trials: Cognitive interview findings from the Pediatric Patient-Reported Outcomes version of the Common Terminology Criteria for Adverse Events initiative. Pediatric blood & cancer. 2017;64(3). |  |  |  |
| RG299 | Reeves BC, Gaus W. Guidelines for reporting non-randomised studies. Forschende Komplementarmedizin und klassische Naturheilkunde = Research in complementary and natural classical medicine. 2004;11 Suppl 1:46-52. |  |  |  |
| RG300 | Ridgewell E, Dobson F, Bach T, Baker R. A systematic review to determine best practice reporting guidelines for AFO interventions in studies involving children with cerebral palsy. Prosthetics and orthotics international. 2010;34(2):129-45. |  |  |  |
| RG301 | Riley RD, Lambert PC, Abo-Zaid G. Meta-analysis of individual participant data: rationale, conduct, and reporting. BMJ (Clinical research ed). 2010;340:c221. |  |  |  |
| RG302 | Robb SL, Burns DS, Carpenter JS. Reporting guidelines for music-based interventions. Journal of health psychology. 2011;16(2):342-52. |  |  |  |
| RG303 | Robinet P, Milewicz DM, Cassis LA, Leeper NJ, Lu HS, Smith JD. Consideration of Sex Differences in Design and Reporting of Experimental Arterial Pathology Studies-Statement From ATVB Council. Arteriosclerosis, thrombosis, and vascular biology. 2018;38(2):292-303. |  |  |  |
| RG304 | Rochon PA, Hoey J, Chan AW, Ferris LE, Lexchin J, Kalkar SR, et al. Financial Conflicts of Interest Checklist 2010 for clinical research studies. Open medicine : a peer-reviewed, independent, open-access journal. 2010;4(1):e69-91. |  |  |  |
| RG305 | Rodgers M, Thomas S, Harden M, Parker G, Street A, Eastwood A. Developing a methodological framework for organisational case studies: a rapid review and consensus development process. Health Serv Deliv Res. 2016;4(1). |  |  |  |
| RG306 | Rosa N. Standards for reporting results of refractive surgery. Journal of refractive surgery (Thorofare, NJ : 1995). 2001;17(4):473-4. |  |  |  |
| RG307 | Rosenthal R, Hoffmann H, Clavien PA, Bucher HC, Dell-Kuster S. Definition and Classification of Intraoperative Complications (CLASSIC): Delphi Study and Pilot Evaluation. World journal of surgery. 2015;39(7):1663-71. | CLASSIC |  |  |
| RG308 | Rothman DJ, McDonald WJ, Berkowitz CD, Chimonas SC, DeAngelis CD, Hale RW, et al. Professional medical associations and their relationships with industry: a proposal for controlling conflict of interest. Jama. 2009;301(13):1367-72. |  |  |  |
| RG309 | Rotta I, Salgado TM, Felix DC, Souza TT, Correr CJ, Fernandez-Llimos F. Ensuring consistent reporting of clinical pharmacy services to enhance reproducibility in practice: an improved version of DEPICT. Journal of evaluation in clinical practice. 2015;21(4):584-90. | DEPICT 2 |  |  |
| RG310 | Rubino M, Pragnell M. Guidelines for reporting case series of tumours of the colon and rectum. Techniques in Coloproctology. 1999;3(2):93-7. |  |  |  |
| RG311 | Rundback JH, Sacks D, Kent KC, Cooper C, Jones D, Murphy T, et al. Guidelines for the reporting of renal artery revascularization in clinical trials. American Heart Association. Circulation. 2002;106(12):1572-85. |  |  |  |
| RG312 | Sacks D, Marinelli DL, Martin LG, Spies JB. Reporting standards for clinical evaluation of new peripheral arterial revascularization devices. Journal of vascular and interventional radiology : JVIR. 2003;14(9 Pt 2):S395-404. |  |  |  |
| RG313 | Saint-Raymond A, Hill S, Martines J, Bahl R, Fontaine O, Bero L. CONSORT 2010 (CONSORT-C). Lancet (London, England). 2010;376(9737):229-30. | CONSORT-C(hildren) |  |  |
| RG314 | Saito M, Gilder ME, Nosten F, Guérin PJ, McGready R. Methodology of assessment and reporting of safety in anti-malarial treatment efficacy studies of uncomplicated falciparum malaria in pregnancy: a systematic literature review. Malaria Journal. 2017;16(1):491. |  |  |  |
| RG315 | Salem R, Lewandowski RJ, Gates VL, Nutting CW, Murthy R, Rose SC, et al. Research reporting standards for radioembolization of hepatic malignancies. Journal of vascular and interventional radiology : JVIR. 2011;22(3):265-78. |  |  |  |
| RG316 | Sanders GD, Neumann PJ, Basu A, Brock DW, Feeny D, Krahn M, et al. Recommendations for Conduct, Methodological Practices, and Reporting of Cost-effectiveness Analyses: Second Panel on Cost-Effectiveness in Health and Medicine. Jama. 2016;316(10):1093-103. |  |  |  |
| RG317 | Sargeant JM, O'Connor AM, Dohoo IR, Erb HN, Cevallos M, Egger M, et al. Methods and processes of developing the strengthening the reporting of observational studies in epidemiology - veterinary (STROBE-Vet) statement. Preventive veterinary medicine. 2016;134:188-96. | STROBE-Vet |  |  |
| RG318 | Scher HI, Eisenberger M, D'Amico AV, Halabi S, Small EJ, Morris M, et al. Eligibility and outcomes reporting guidelines for clinical trials for patients in the state of a rising prostate-specific antigen: recommendations from the Prostate-Specific Antigen Working Group. Journal of clinical oncology: official journal of the American Society of Clinical Oncology. 2004;22(3):537-56. |  |  |  |
| RG319 | Schreiber JB. Latent Class Analysis: An example for reporting results. Research in Social and Administrative Pharmacy. 2017;13(6):1196-201. |  |  |  |
| RG320 | Schriger DL. Suggestions for Improving the Reporting of Clinical Research: The Role of Narrative. Annals of Emergency Medicine. 2005;45(4):437-43. |  |  |  |
| RG321 | Schulz KF, Altman DG, Moher D. CONSORT 2010 statement: updated guidelines for reporting parallel group randomised trials. BMJ (Clinical research ed). 2010;340:c332. | CONSORT |  |  |
| RG322 | Sechopoulos I, Rogers DWO, Bazalova-Carter M, Bolch WE, Heath EC, McNitt-Gray MF, et al. RECORDS: improved Reporting of montE CarlO RaDiation transport Studies: Report of the AAPM Research Committee Task Group 268. Medical physics. 2018;45(1):e1-e5. | RECORDS |  |  |
| RG323 | Sena ES, Currie GL, McCann SK, Macleod MR, Howells DW. Systematic reviews and meta-analysis of preclinical studies: why perform them and how to appraise them critically. Journal of cerebral blood flow and metabolism : official journal of the International Society of Cerebral Blood Flow and Metabolism. 2014;34(5):737-42. |  |  |  |
| RG324 | Sepucha KR, Abhyankar P, Hoffman AS, Bekker HL, LeBlanc A, Levin CA, et al. Standards for UNiversal reporting of patient Decision Aid Evaluation studies: the development of SUNDAE Checklist. BMJ quality & safety. 2018;27(5):380-8. | SUNDAE |  |  |
| RG325 | Sharp SJ, Poulaliou M, Thompson SG, White IR, Wood AM. A review of published analyses of case-cohort studies and recommendations for future reporting. PloS one. 2014;9(6):e101176-e. |  |  |  |
| RG326 | Shemin RJ, Cox JL, Gillinov AM, Blackstone EH, Bridges CR. Guidelines for reporting data and outcomes for the surgical treatment of atrial fibrillation. The Annals of thoracic surgery. 2007;83(3):1225-30. |  |  |  |
| RG327 | Shiffman RN, Shekelle P, Overhage JM, Slutsky J, Grimshaw J, Deshpande AM. Standardized reporting of clinical practice guidelines: a proposal from the Conference on Guideline Standardization. Annals of internal medicine. 2003;139(6):493-8. |  |  |  |
| RG328 | Simel DL, Rennie D, Bossuyt PM. The STARD statement for reporting diagnostic accuracy studies: application to the history and physical examination. Journal of general internal medicine. 2008;23(6):768-74. |  |  |  |
| RG329 | Singh JP, Yang S, Mulvey EP. Reporting guidance for violence risk assessment predictive validity studies: the RAGEE Statement. Law and human behavior. 2015;39(1):15-22. |  |  |  |
| RG330 | Siontis GC, Patsopoulos NA, Vlahos AP, Ioannidis JP. Selection and presentation of imaging figures in the medical literature. PloS one. 2010;5(5):e10888. |  |  |  |
| RG331 | Slade SC, Dionne CE, Underwood M, Buchbinder R, Beck B, Bennell K, et al. Consensus on Exercise Reporting Template (CERT): Modified Delphi Study. Physical therapy. 2016;96(10):1514-24. | CERT |  |  |
| RG332 | Slemc L, Kunej T. Transcription factor HIF1A: downstream targets, associated pathways, polymorphic hypoxia response element (HRE) sites, and initiative for standardization of reporting in scientific literature. Tumour biology : the journal of the International Society for Oncodevelopmental Biology and Medicine. 2016;37(11):14851-61. |  |  |  |
| RG333 | Smith JA, Arshad Z, Trippe A, Collins GS, Brindley DA, Carr AJ. The Reporting Items for Patent Landscapes statement. Nature biotechnology. 2018;36(11):1043-7. | RIPL |  |  |
| RG334 | Smith L, Rosenzweig L, Schmidt M. Best Practices in the Reporting of Participatory Action Research: Embracing Both the Forest and the Trees. The Counseling Psychologist. 2010;38(8):1115-38. |  |  |  |
| RG335 | Smith SM, Hunsinger M, McKeown A, Parkhurst M, Allen R, Kopko S, et al. Quality of pain intensity assessment reporting: ACTTION systematic review and recommendations. The journal of pain : official journal of the American Pain Society. 2015;16(4):299-305. |  |  |  |
| RG336 | Soong CV, Dasari BV, Loan W, Hannon R, Lee B, Lau L, et al. Setting the standards for reporting ruptured abdominal aortic aneurysm. Vascular and endovascular surgery. 2010;44(6):449-53. |  |  |  |
| RG337 | Sorinola O, Olufowobi O, Coomarasamy A, Khan KS. Instructions to authors for case reporting are limited: a review of a core journal list. BMC medical education. 2004;4:4-. |  |  |  |
| RG338 | Spiegelhalter DJ, Myles JP, Jones DR, Abrams KR. Bayesian methods in health technology assessment: a review. Health technology assessment (Winchester, England). 2000;4(38):1-130. |  |  |  |
| RG339 | Stamp LK, Morillon MB, Taylor WJ, Dalbeth N, Singh JA, Lassere M, et al. Variability in the Reporting of Serum Urate and Flares in Gout Clinical Trials: Need for Minimum Reporting Requirements. The Journal of rheumatology. 2018;45(3):419-24. |  |  |  |
| RG340 | Staniszewska S, Brett J, Simera I, Seers K, Mockford C, Goodlad S, et al. GRIPP2 reporting checklists: tools to improve reporting of patient and public involvement in research. BMJ (Clinical research ed). 2017;358:j3453. | GRIPP2 |  |  |
| RG341 | Staquet M, Berzon R, Osoba D, Machin D. Guidelines for reporting results of quality of life assessments in clinical trials. Quality of life research : an international journal of quality of life aspects of treatment, care and rehabilitation. 1996;5(5):496-502. |  |  |  |
| RG342 | Steele J, Fisher J, Giessing J, Gentil P. Clarity in reporting terminology and definitions of set endpoints in resistance training. Muscle & nerve. 2017;56(3):368-74. |  |  |  |
| RG343 | Sterne JA, White IR, Carlin JB, Spratt M, Royston P, Kenward MG, et al. Multiple imputation for missing data in epidemiological and clinical research: potential and pitfalls. BMJ (Clinical research ed). 2009;338:b2393. |  |  |  |
| RG344 | Stevens GA, Alkema L, Black RE, Boerma JT, Collins GS, Ezzati M, et al. Guidelines for Accurate and Transparent Health Estimates Reporting: the GATHER statement. Lancet (London, England). 2016;388(10062):e19-e23. | GATHER |  |  |
| RG345 | Stewart LA, Clarke M, Rovers M, Riley RD, Simmonds M, Stewart G, et al. Preferred Reporting Items for Systematic Review and Meta-Analyses of individual participant data: the PRISMA-IPD Statement. Jama. 2015;313(16):1657-65. | PRISMA-IPD |  |  |
| RG346 | Stiles CR, Biondo PD, Cummings G, Hagen NA. Clinical trials focusing on cancer pain educational interventions: core components to include during planning and reporting. Journal of pain and symptom management. 2010;40(2):301-8. |  |  |  |
| RG347 | Stock-Schroer B, Albrecht H, Betti L, Endler PC, Linde K, Ludtke R, et al. Reporting experiments in homeopathic basic research (REHBaR)--a detailed guideline for authors. Homeopathy : the journal of the Faculty of Homeopathy. 2009;98(4):287-98. | REHBaR |  |  |
| RG348 | Stone AA, Shiffman S. Capturing momentary, self-report data: a proposal for reporting guidelines. Annals of behavioral medicine : a publication of the Society of Behavioral Medicine. 2002;24(3):236-43. |  |  |  |
| RG349 | Stone SP, Cooper BS, Kibbler CC, Cookson BD, Roberts JA, Medley GF, et al. The ORION statement: guidelines for transparent reporting of outbreak reports and intervention studies of nosocomial infection. The Lancet Infectious diseases. 2007;7(4):282-8. | ORION |  |  |
| RG350 | Stout NK, Knudsen AB, Kong CY, McMahon PM, Gazelle GS. Calibration methods used in cancer simulation models and suggested reporting guidelines. PharmacoEconomics. 2009;27(7):533-45. |  |  |  |
| RG351 | Stroup DF, Berlin JA, Morton SC, Olkin I, Williamson GD, Rennie D, et al. Meta-analysis of observational studies in epidemiology: a proposal for reporting. Meta-analysis Of Observational Studies in Epidemiology (MOOSE) group. Jama. 2000;283(15):2008-12. | MOOSE |  |  |
| RG352 | Subramanian J, Simon R. Gene expression-based prognostic signatures in lung cancer: ready for clinical use? Journal of the National Cancer Institute. 2010;102(7):464-74. |  |  |  |
| RG353 | Sun BC, Thiruganasambandamoorthy V, Cruz JD. Standardized reporting guidelines for emergency department syncope risk-stratification research. Academic emergency medicine : official journal of the Society for Academic Emergency Medicine. 2012;19(6):694-702. |  |  |  |
| RG354 | Sung L, Hayden J, Greenberg ML, Koren G, Feldman BM, Tomlinson GA. Seven items were identified for inclusion when reporting a Bayesian analysis of a clinical study. Journal of clinical epidemiology. 2005;58(3):261-8. |  |  |  |
| RG355 | Systematic Reviews. CRD's guidance for undertaking reviews in health care. Centre for Reviews and Dissemination, University of York; 2009. https://www.york.ac.uk/media/crd/Systematic_Reviews.pdf |  |  |  |
| RG356 | Tacconelli E, Cataldo MA, Paul M, Leibovici L, Kluytmans J, Schroder W, et al. STROBE-AMS: recommendations to optimise reporting of epidemiological studies on antimicrobial resistance and informing improvement in antimicrobial stewardship. BMJ Open. 2016;6(2):e010134. | STROBE-AMS |  |  |
| RG357 | Talmon J, Ammenwerth E, Brender J, de Keizer N, Nykanen P, Rigby M. STARE-HI--Statement on reporting of evaluation studies in Health Informatics. International journal of medical informatics. 2009;78(1):1-9. |  |  |  |
| RG358 | Tate RL, Perdices M, Rosenkoetter U, Shadish W, Vohra S, Barlow DH, et al. The Single-Case Reporting Guideline In BEhavioural Interventions (SCRIBE) 2016 Statement. Physical therapy. 2016;96(7):e1-e10. | SCRIBE |  |  |
| RG359 | ter Haar G, Shaw A, Pye S, Ward B, Bottomley F, Nolan R, et al. Guidance on reporting ultrasound exposure conditions for bio-effects studies. Ultrasound in medicine & biology. 2011;37(2):177-83. |  |  |  |
| RG360 | Timaran CH, McKinsey JF, Schneider PA, Littooy F. Reporting standards for carotid interventions from the Society for Vascular Surgery. Journal of vascular surgery. 2011;53(6):1679-95. |  |  |  |
| RG361 | Tomaszewski KA, Henry BM, Kumar Ramakrishnan P, Roy J, Vikse J, Loukas M, et al. Development of the Anatomical Quality Assurance (AQUA) checklist: Guidelines for reporting original anatomical studies. Clinical anatomy (New York, NY). 2017;30(1):14-20. | AQUA |  |  |
| RG362 | Tong A, Flemming K, McInnes E, Oliver S, Craig J. Enhancing transparency in reporting the synthesis of qualitative research: ENTREQ. BMC Med Res Methodol. 2012;12:181. | ENTREQ |  |  |
| RG363 | Tong A, Sainsbury P, Craig J. Consolidated criteria for reporting qualitative research (COREQ): a 32-item checklist for interviews and focus groups. International journal for quality in health care : journal of the International Society for Quality in Health Care. 2007;19(6):349-57. | COREQ |  |  |
| RG364 | Tricco AC, Lillie E, Zarin W, O'Brien KK, Colquhoun H, Levac D, et al. PRISMA Extension for Scoping Reviews (PRISMA-ScR): Checklist and Explanation. Annals of internal medicine. 2018;169(7):467-73. | PRISMA-ScR |  |  |
| RG365 | Turina MI, Shennib H, Dunning J, Cheng D, Martin J, Muneretto C, et al. EACTS/ESCVS best practice guidelines for reporting treatment results in the thoracic aorta. European journal of cardio-thoracic surgery : official journal of the European Association for Cardio-thoracic Surgery. 2009;35(6):927-30. |  |  |  |
| RG366 | van de Schoot R, Sijbrandij M, Winter SD, Depaoli S, Vermunt JK. The GRoLTS-Checklist: Guidelines for Reporting on Latent Trajectory Studies. Structural Equation Modeling: A Multidisciplinary Journal. 2017;24(3):451-67. | GRoLTS |  |  |
| RG367 | van Haselen RA. Homeopathic clinical case reports: Development of a supplement (HOM-CASE) to the CARE clinical case reporting guideline. Complementary therapies in medicine. 2016;25:78-85. | HOM-CASE |  |  |
| RG368 | Vanhie A, Meuleman C, Tomassetti C, Timmerman D, D'Hoore A, Wolthuis A, et al. Consensus on Recording Deep Endometriosis Surgery: the CORDES statement. Human reproduction (Oxford, England). 2016;31(6):1219-23. | CORDES |  |  |
| RG369 | Vasilevsky NA, Brush MH, Paddock H, Ponting L, Tripathy SJ, Larocca GM, et al. On the reproducibility of science: unique identification of research resources in the biomedical literature. PeerJ. 2013;1:e148. |  |  |  |
| RG370 | Vasquez MA, Munschauer CE. The importance of uniform venous terminology in reports on varicose veins. Seminars in vascular surgery. 2010;23(2):70-7. |  |  |  |
| RG371 | Vernooij RWM, Alonso-Coello P, Brouwers M, Martínez García L, CheckUp P. Reporting Items for Updated Clinical Guidelines: Checklist for the Reporting of Updated Guidelines (CheckUp). PLoS medicine. 2017;14(1):e1002207-e. | CheckUp |  |  |
| RG372 | Vickers AJ, Ballen V, Scher HI. Setting the bar in phase II trials: the use of historical data for determining "go/no go" decision for definitive phase III testing. Clinical cancer research: an official journal of the American Association for Cancer Research. 2007;13(3):972-6. |  |  |  |
| RG373 | Vijayakumar N, Mills KL, Alexander-Bloch A, Tamnes CK, Whittle S. Structural brain development: A review of methodological approaches and best practices. Developmental cognitive neuroscience. 2018;33:129-48. |  |  |  |
| RG374 | Vintzileos AM, Beazoglou T. Design, execution, interpretation, and reporting of economic evaluation studies in obstetrics. American journal of obstetrics and gynecology. 2004;191(4):1070-6. |  |  |  |
| RG375 | Virues-Ortega J, Moreno-Rodriguez R. Guidelines for clinical case reports in behavioral clinical psychology. Int J Clin Health Psychology. 2008;8(3):765-77. |  |  |  |
| RG376 | Vitek JL, Lyons KE, Bakay R, Benabid AL, Deuschl G, Hallett M, et al. Standard guidelines for publication of deep brain stimulation studies in Parkinson's disease (Guide4DBS-PD). Movement disorders: official journal of the Movement Disorder Society. 2010;25(11):1530-7. | Guide4DBS-PD |  |  |
| RG377 | Vohra S, Shamseer L, Sampson M, Bukutu C, Schmid CH, Tate R, et al. CONSORT extension for reporting N-of-1 trials (CENT) 2015 Statement. Journal of clinical epidemiology. 2016;76:9-17. | CONSORT-CENT |  |  |
| RG378 | von Elm E, Altman DG, Egger M, Pocock SJ, Gotzsche PC, Vandenbroucke JP. Strengthening the Reporting of Observational Studies in Epidemiology (STROBE) statement: guidelines for reporting observational studies. BMJ (Clinical research ed). 2007;335(7624):806-8. | STROBE |  |  |
| RG379 | Walker MF, Hoffmann TC, Brady MC, Dean CM, Eng JJ, Farrin AJ, et al. Improving the development, monitoring and reporting of stroke rehabilitation research: Consensus-based core recommendations from the Stroke Recovery and Rehabilitation Roundtable. International journal of stroke : official journal of the International Stroke Society. 2017;12(5):472-9. |  |  |  |
| RG380 | Wang R, Lagakos SW, Ware JH, Hunter DJ, Drazen JM. Statistics in medicine--reporting of subgroup analyses in clinical trials. The New England journal of medicine. 2007;357(21):2189-94. |  |  |  |
| RG381 | Wang SV, Schneeweiss S, Berger ML, Brown J, de Vries F, Douglas I, et al. Reporting to Improve Reproducibility and Facilitate Validity Assessment for Healthcare Database Studies V1.0. Pharmacoepidemiology and drug safety. 2017;26(9):1018-32. |  |  |  |
| RG382 | Wardlaw JM, Smith EE, Biessels GJ, Cordonnier C, Fazekas F, Frayne R, et al. Neuroimaging standards for research into small vessel disease and its contribution to ageing and neurodegeneration. The Lancet Neurology. 2013;12(8):822-38. |  |  |  |
| RG383 | Webster JD, Dennis MM, Dervisis N, Heller J, Bacon NJ, Bergman PJ, et al. Recommended guidelines for the conduct and evaluation of prognostic studies in veterinary oncology. Veterinary pathology. 2011;48(1):7-18. |  |  |  |
| RG384 | Welch RW, Antoine JM, Berta JL, Bub A, de Vries J, Guarner F, et al. Guidelines for the design, conduct and reporting of human intervention studies to evaluate the health benefits of foods. The British journal of nutrition. 2011;106 Suppl 2:S3-15. |  |  |  |
| RG385 | Welch V, Petticrew M, Tugwell P, Moher D, O'Neill J, Waters E, et al. PRISMA-Equity 2012 extension: reporting guidelines for systematic reviews with a focus on health equity. PLoS medicine. 2012;9(10):e1001333. | PRISMA-Equity 2012 |  |  |
| RG386 | Welch VA, Norheim OF, Jull J, Cookson R, Sommerfelt H, Tugwell P. CONSORT-Equity 2017 extension and elaboration for better reporting of health equity in randomised trials. BMJ (Clinical research ed). 2017;359:j5085. | CONSORT-Equity |  |  |
| RG387 | Wendler JJ, Fischbach K, Ricke J, Jurgens J, Fischbach F, Kollermann J, et al. Irreversible Electroporation (IRE): Standardization of Terminology and Reporting Criteria for Analysis and Comparison. Polish journal of radiology. 2016;81:54-64. |  |  |  |
| RG388 | White A. Conducting and reporting case series and audits--author guidelines for acupuncture in medicine. Acupuncture in medicine : journal of the British Medical Acupuncture Society. 2005;23(4):181-7. |  |  |  |
| RG389 | White RG, Hakim AJ, Salganik MJ, Spiller MW, Johnston LG, Kerr L, et al. Strengthening the Reporting of Observational Studies in Epidemiology for respondent-driven sampling studies: "STROBE-RDS" statement. Journal of clinical epidemiology. 2015;68(12):1463-71. | STROBE-RDS |  |  |
| RG390 | Widmann G, Stoffner R, Sieb M, Bale R. Target registration and target positioning errors in computer-assisted neurosurgery: proposal for a standardized reporting of error assessment. The international journal of medical robotics + computer assisted surgery : MRCAS. 2009;5(4):355-65. |  |  |  |
| RG391 | Winchester C. AMWA‒EMWA‒ISMPP Joint Position Statement on the Role of Professional Medical Writers. Medical Writing. 2017;26(1). |  |  |  |
| RG392 | Witkiewitz K, Finney JW, Harris AH, Kivlahan DR, Kranzler HR. Guidelines for the Reporting of Treatment Trials for Alcohol Use Disorders. Alcoholism, clinical and experimental research. 2015;39(9):1571-81. |  |  |  |
| RG393 | Wolfe F, Lassere M, van der Heijde D, Stucki G, Suarez-Almazor M, Pincus T, et al. Preliminary core set of domains and reporting requirements for longitudinal observational studies in rheumatology. The Journal of rheumatology. 1999;26(2):484-9. |  |  |  |
| RG394 | Wong G, Greenhalgh T, Westhorp G, Buckingham J, Pawson R. RAMESES publication standards: meta-narrative reviews. BMC medicine. 2013;11:20. RAMESES: meta-narrative reviews | RAMESES: meta-narrative reviews |  |  |
| RG395 | Wong G, Greenhalgh T, Westhorp G, Buckingham J, Pawson R. RAMESES publication standards: realist syntheses. BMC medicine. 2013;11:21. | RAMESES: realist syntheses |  |  |
| RG396 | Wong G, Westhorp G, Manzano A, Greenhalgh J, Jagosh J, Greenhalgh T. RAMESES II reporting standards for realist evaluations. BMC medicine. 2016;14(1):96. | RAMESES II |  |  |
| RG397 | Wu T, Shang H, Bian Z, Zhang J, Li T, Li Y, et al. Recommendations for reporting adverse drug reactions and adverse events of traditional Chinese medicine. Journal of evidence-based medicine. 2010;3(1):11-7. |  |  |  |
| RG398 | Xie F, Pickard AS, Krabbe PF, Revicki D, Viney R, Devlin N, et al. A Checklist for Reporting Valuation Studies of Multi-Attribute Utility-Based Instruments (CREATE). PharmacoEconomics. 2015;33(8):867-77. | CREATE |  |  |
| RG399 | Yao XI, Wang X, Speicher PJ, Hwang ES, Cheng P, Harpole DH, et al. Reporting and Guidelines in Propensity Score Analysis: A Systematic Review of Cancer and Cancer Surgical Studies. Journal of the National Cancer Institute. 2017;109(8). |  |  |  |
| RG400 | Yong MY, Gonzalez-Beltran A, Begent R. Establishing a knowledge trail from molecular experiments to clinical trials. New biotechnology. 2011;28(5):464-80. |  |  |  |
| RG401 | Yousafzai AK, Aboud FE, Nores M, Kaur R. Reporting guidelines for implementation research on nurturing care interventions designed to promote early childhood development. Annals of the New York Academy of Sciences. 2018;1419(1):26-37. | C.A.R.E. |  |  |
| RG402 | Zakrzewska JM, Lopez BC. Quality of reporting in evaluations of surgical treatment of trigeminal neuralgia: recommendations for future reports. Neurosurgery. 2003;53(1):110-20; discussion 20-2. |  |  |  |
| RG403 | Zanoni G, Girolomoni G, Bonetto C, Trotta F, Hausermann P, Opri R, et al. Single organ cutaneous vasculitis: Case definition & guidelines for data collection, analysis, and presentation of immunization safety data. Vaccine. 2016;34(51):6561-71. |  |  |  |
| RG404 | Zaritsky A, Nadkarni V, Hazinski MF, Foltin G, Quan L, Wright J, et al. Recommended guidelines for uniform reporting of pediatric advanced life support: the Pediatric Utstein Style. A statement for healthcare professionals from a task force of the American Academy of Pediatrics, the American Heart Association, and the European Resuscitation Council. Resuscitation. 1995;30(2):95-115. |  |  |  |
| RG405 | Zavada J, Dixon WG, Askling J. Launch of a checklist for reporting longitudinal observational drug studies in rheumatology: a EULAR extension of STROBE guidelines based on experience from biologics registries. Annals of the rheumatic diseases. 2014;73(3):628. |  |  |  |
| RG406 | Zorzela L, Loke YK, Ioannidis JP, Golder S, Santaguida P, Altman DG, et al. PRISMA harms checklist: improving harms reporting in systematic reviews. BMJ (Clinical research ed). 2016;352:i157. | PRISMA harms |  |  |
| RG407 | Zwarenstein M, Treweek S, Gagnier JJ, Altman DG, Tunis S, Haynes B, et al. Improving the reporting of pragmatic trials: an extension of the CONSORT statement. BMJ (Clinical research ed). 2008;337:a2390. | CONSORT Pragmatic trials |  |  |

S2 Table. Comparison of electronic and manual identification of sex and gender related words in reporting guidelines

| Id | Checklist | | | | | | Statement | | | | | | | | | | References | | | | | | | |
| --- | --- | --- | --- | --- | --- | --- | --- | --- | --- | --- | --- | --- | --- | --- | --- | --- | --- | --- | --- | --- | --- | --- | --- | --- |
|  | **Sex** | | **Men** | | **Women** | | **Gender** | | **Male** | | **Woman** | | **Men** | | **Women** | | **Sex** | | **Gender** | | **Men** | | **Women** | |
|  | **M** | **E** | **M** | **E** | **M** | **E** | **M** | **E** | **M** | **E** | **M** | **E** | **M** | **E** | **M** | **E** | **M** | **E** | **M** | **E** | **M** | **E** | **M** | **E** |
| 37 |  |  | 0 | **1** | 0 | **1** |  |  |  |  |  |  |  |  |  |  |  |  |  |  |  |  |  |  |
| 50 |  |  |  |  |  |  |  |  |  |  |  |  | 0 | **3** | 0 | **4** |  |  |  |  |  |  |  |  |
| 60 |  |  |  |  |  |  |  |  |  |  |  |  | 14 | **15** |  |  |  |  |  |  |  |  |  |  |
| 79 |  |  |  |  |  |  | 0 | **1** |  |  |  |  |  |  |  |  |  |  |  |  |  |  |  |  |
| 123 |  |  |  |  |  |  |  |  |  |  |  |  |  |  |  |  | 0 | **1** |  |  |  |  |  |  |
| 129 |  |  |  |  |  |  |  |  |  |  | 0 | 2 |  |  |  |  |  |  |  |  |  |  |  |  |
| 189 |  |  |  |  |  |  |  |  | 3 | **4** |  |  |  |  |  |  |  |  |  |  |  |  |  |  |
| 263 |  |  |  |  |  |  | 1 | **3** |  |  |  |  | 0 | **2** |  |  |  |  | **7** | 6 |  |  |  |  |
| 303 |  |  |  |  |  |  |  |  |  |  |  |  |  |  |  |  |  |  |  |  | 0 | **1** | 0 | **1** |
| 311 | 0 | **1** |  |  |  |  |  |  |  |  |  |  |  |  |  |  |  |  |  |  |  |  |  |  |
| 323 |  |  |  |  |  |  |  |  |  |  |  |  |  |  |  |  | 0 | **1** |  |  |  |  |  |  |
| 385 |  |  |  |  |  |  |  |  |  |  |  |  |  |  | 4 | **2** |  |  |  |  |  |  |  |  |

E: Electronic search; M : manual search; correct value in green after verification

S3 Table. Concordance between electronic and manual identification of sex and gender related words in reporting guidelines

|  | **Electronic** | | | | **Manual** | | | |  |
| --- | --- | --- | --- | --- | --- | --- | --- | --- | --- |
| **Characteristics** | | | | | | | | | |
|  | Min_e_ | Max_e_ | Mean_e_ | Med_e_ | Min_m_ | Max_m_ | Mean_m_ | Med_m_ | Conc % |
| **Checklist** | | | | | | | | | |
| Sex | 0 | 1 | 0.02439 | 0 | 0 | 0 | 0 | 0 | 97.6 |
| Gender | 0 | 3 | 0.12195 | 0 | 0 | 3 | 0.12195 | 0 | 100 |
| Woman | 0 | 0 | 0 | 0 | 0 | 0 | 0 | 0 | 100 |
| Women | 0 | 1.0000 | 0.02439 | 0 | 0 | 0 | 0 | 0 | 97.6 |
| Man | 0 | 0 | 0 | 0 | 0 | 0 | 0 | 0 | 100 |
| Men | 0 | 1.0000 | 0.02439 | 0 | 0 | 0 | 0 | 0 | 97.6 |
| Female | 0 | 1.0000 | 0.04878 | 0 | 0 | 1.0000 | 0.04878 | 0 | 100 |
| Girl | 0 | 0 | 0 | 0 | 0 | 0 | 0 | 0 | 100 |
| Boy | 0 | 0 | 0 | 0 | 0 | 0 | 0 | 0 | 100 |
| Male | 0 | 1.0000 | 0.04878 | 0 | 0 | 1.0000 | 0.04878 | 0 | 100 |
| Total |  | | | | | | | | 99.3 |
| **Abstract** | | | | | | | | | |
| Sex | 0 | 0 | 0 | 0 | 0 | 0 | 0 | 0 | 100 |
| Gender | 0 | 0 | 0 | 0 | 0 | 0 | 0 | 0 | 100 |
| Woman | 0 | 0 | 0 | 0 | 0 | 0 | 0 | 0 | 100 |
| Women | 0 | 0 | 0 | 0 | 0 | 0 | 0 | 0 | 100 |
| Man | 0 | 0 | 0 | 0 | 0 | 0 | 0 | 0 | 100 |
| Men | 0 | 0 | 0 | 0 | 0 | 0 | 0 | 0 | 100 |
| Female | 0 | 0 | 0 | 0 | 0 | 0 | 0 | 0 | 100 |
| Girl | 0 | 0 | 0 | 0 | 0 | 0 | 0 | 0 | 100 |
| Boy | 0 | 0 | 0 | 0 | 0 | 0 | 0 | 0 | 100 |
| Male | 0 | 0 | 0 | 0 | 0 | 0 | 0 | 0 | 100 |
| Total |  | | | | | | | | 100 |
| **Statement** | | | | | | | | | |
| Sex | 0 | 30 | 0.90243 | 0 | 0 | 30 | 0.90243 | 0 | 100 |
| Gender | 0 | 4 | 0.34146 | 0 | 0 | 4 | 0.26829 | 0 | 95.1 |
| Woman | 0 | 2 | 0.04878 | 0 | 0 | 0 | 0 | 0 | 97.6 |
| Women | 0 | 17 | 0.75609 | 0 | 0 | 17 | 0.70731 | 0 | 95.1 |
| Man | 0 | 0 | 0 | 0 | 0 | 0 | 0 | 0 | 100 |
| Men | 0 | 15 | 0.56097 | 0 | 0 | 14 | 0.41463 | 0 | 95.1 |
| Female | 0 | 2 | 0.07317 | 0 | 0 | 2 | 0.07317 | 0 | 100 |
| Girl | 0 | 0 | 0 | 0 | 0 | 0 | 0 | 0 | 100 |
| Boy | 0 | 0 | 0 | 0 | 0 | 0 | 0 | 0 | 100 |
| Male | 0 | 4 | 0.14634 | 0 | 0 | 3 | 0.12195 | 0 | 97.6 |
| Total |  | | | | | | | | 98.1 |
| **References** | | | | | | | | | |
| Sex | 0 | 1 | 0.12195 | 0 | 0 | 1 | 0.07317 | 0 | 95.1 |
| Gender | 0 | 6 | 0.14634 | 0 | 0 | 7 | 0.17073 | 0 | 97.6 |
| Woman | 0 | 0 | 0 | 0 | 0 | 0 | 0 | 0 | 100 |
| Women | 0 | 9 | 0.43902 | 0 | 0 | 9 | 0.41463 | 0 | 97.6 |
| Man | 0 | 0 | 0 | 0 | 0 | 0 | 0 | 0 | 100 |
| Men | 0 | 2 | 0.09756 | 0 | 0 | 2 | 0.07317 | 0 | 97.6 |
| Female | 0 | 0 | 0 | 0 | 0 | 0 | 0 | 0 | 100 |
| Girl | 0 | 0 | 0 | 0 | 0 | 0 | 0 | 0 | 100 |
| Boy | 0 | 1 | 0.02439 | 0 | 0 | 1 | 0.02439 | 0 | 100 |
| Male | 0 | 0 | 0 | 0 | 0 | 0 | 0 | 0 | 100 |
| Total |  |  |  |  |  |  |  |  | 98.8 |

Min : minimum ; Max : maximum ; Med : median ; Conc: concordance

S4 Table. Distribution of the use of “sex” in various study types and sections of reporting guidelines

|  | **Section of reporting guideline** | | | |  |
| --- | --- | --- | --- | --- | --- |
|  | Checklist | Flowchart | Abstract | Statement | All^1^ |
| **Study type** | | | | |  |
| Case report^2^ | 1 (25) | 0 (0) | 0 (0) | 0 (0) | 1 (25) |
| Clinical practice guideline | 0 (0) | 0 (0) | 0 (0) | 0 (0) | 0 (0) |
| Diagnostic/prognostic | 8 (47.6) | 0 (0) | 0 (0) | 2 (11.1) | 9 (50) |
| Economic evaluation | 0 (0) | 0 (0) | 0 (0) | 2 (12.5) | 3 (18.8) |
| Experiment | 11 (8.6) | 0 (0) | 0 (0) | 18 (12.3) | 30 (29.4) |
| Nonspecific^3^ | 12 (14.5) | 1 (25) | 1 (1.3) | 15 (14.6) | 24 (23.3) |
| Observational | 21 (18.4) | 0 (0) | 1 (0.9) | 18 (4.4) | 39 (32.5) |
| Other^4^ | 0 (0) | 0 (0) | 0 (0) | 2 (11.8) | 3 (17.7) |
| Preclinical | 5 (1.4) | 0 (0) | 1 (7.7) | 4 (26.7) | 8 (53.3) |
| Protocol | 0 (0) | 0 (0) | 0 (0) | 1 (10) | 1 (10) |
| Quality improvement | 0 (0) | 0 (0) | 0 (0) | 0 (0) | 0 (0) |
| Qualitative | 1 (6.3) | 0 (0) | 0 (0) | 1 (6) | 3 (17.7) |
| Randomised trial | 10 (2.6) | 0 (0) | 0 (0) | 17 (12.8) | 29 (21.8) |
| Systematic review | 5 (1.4) | 0 (0) | 0 (0) | 5 (14.7) | 8 (23.5) |

^1^All sections including references

^2^Based on category of study types of EQUATOR homepage

^3^ Do not apply to any specific type of study

^4^As specified on the individual page of reporting guidelines on EQUATOR

S5 Table. Distribution of the use of “gender” in various study types and sections of reporting guidelines

|  | **Section of reporting guideline** | | | |  |
| --- | --- | --- | --- | --- | --- |
|  | Checklist | Flowchart | Abstract | Statement | All^1^ |
| **Study type** | | | | |  |
| Case report^2^ | 3 (75.0) | 0 (0) | 0 (0) | 0 (0) | 3 (75.0) |
| Clinical practice guideline | 0 (0) | 0 (0) | 0 (0) | 2 (25) | 2 (25) |
| Diagnostic/prognostic | 1 (5.9) | 0 (0) | 0 (0) | 3 (16.7) | 4 (22.2) |
| Economic evaluation | 0 (0) | 0 (0) | 0 (0) | 2 (12.5) | 2 (12.5) |
| Experiment | 9 (7.0) | 0 (0) | 0 (0) | 21 (14.4) | 25 (17.1) |
| Nonspecific^3^ | 9 (2.5) | 1 (25) | 4 (5.2) | 16 (15.5) | 22 (21.4) |
| Observational | 14 (12.3) | 0 (0) | 0 (0) | 26 (21.2) | 31 (25.8) |
| Other^4^ | 2 (12.5) | 0 (0) | 0 (0) | 4 (23.5) | 6 (35.3) |
| Preclinical | 3 (23.1) | 0 (0) | 1 (7.7) | 3 (20) | 5 (33.3) |
| Protocol | 2 (20) | 0 (0) | 0 (0) | 2 (20) | 3 (30) |
| Quality improvement | 0 (0) | 0 (0) | 0 (0) | 1 (20) | 1 (20) |
| Qualitative | 2 (12.5) | 0 (0) | 0 (0) | 4 (23.5) | 4 (23.5) |
| Randomised trial | 7 (6.1) | 0 (0) | 0 (0) | 18 (13.5) | 22 (16.5) |
| Systematic review | 3 (8.8) | 0 (0) | 0 (0) | 7 (20.6) | 9 (26.5) |

^1^All sections including references

^2^Based on category of study types of EQUATOR homepage

^3^ Do not apply to any specific type of study

^4^As specified on the individual page of reporting guidelines on EQUATOR

S1 Image. Example of manual count of sex- and gender-related words

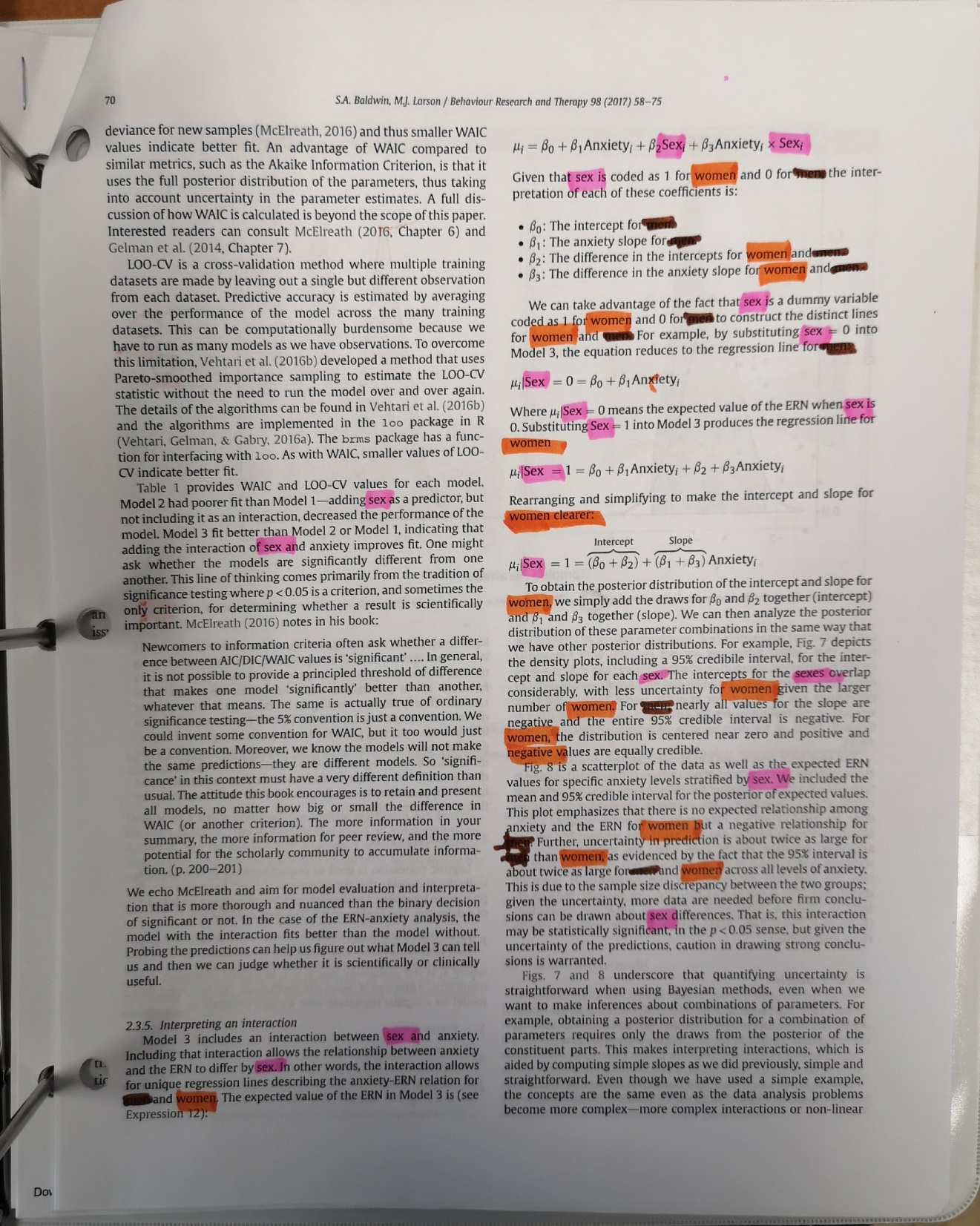
